## supplementary for "Dense, Continuous Membrane Labeling and Expansion Microscopy Visualization of Ultrastructure in Tissues"

### Contents

**Supplementary Notes** ... 3
**Supplementary Note 1** ... 3
**Supplementary Note 2** ... 3
**Supplementary Note 3** ... 4
**Supplementary Note 4** ... 4
**Supplementary Note 5** ... 5
**Supplementary Note 6** ... 5
**Supplementary Figures** ... 6
**Supplementary Figure 1** ... 6
**Supplementary Figure 2** ... 7
**Supplementary Figure 3** ... 9
**Supplementary Figure 4** ... 10
**Supplementary Figure 5** ... 11
**Supplementary Figure 6** ... 12
**Supplementary Figure 7** ... 13
**Supplementary Figure 8** ... 15
**Supplementary Figure 9** ... 16
**Supplementary Figure 10** ... 17
**Supplementary Figure 11** ... 18
**Supplementary Figure 12** ... 19
**Supplementary Figure 13** ... 20
**Supplementary Figure 14** ... 21
**Supplementary Figure 15** ... 23
**Supplementary Figure 16** ... 24
**Supplementary Figure 17** ... 25
**Supplementary Figure 18** ... 26
**Supplementary Figure 19** ... 27
**Supplementary Figure 20** ... 28
**Supplementary Figure 21** ... 29
**Supplementary Figure 22** ... 31

### Supplementary Notes

#### Supplementary Note 1

We incubated mouse brain tissue sections with 100 μM of the membrane probe pGk5a (pGk5b, with an azide replacing the biotin). We post-labeled the specimens with gold nanoparticles modified with a dibenzocyclooctyne (DBCO) handle, for EM visualization (see **Supplementary Methods** for details). We then imaged the resulting specimens with EM. We saw similar details of intact membranes, organelles, and synapses (**Supp. Fig. 1a**), when compared to classical $OsO_4$ membrane visualization (**Supp. Fig. 1b**), in EM.

#### Supplementary Note 2

We evaluated the isotropy of mExM expansion by quantitatively comparing pre-expansion structured illumination microscopy (SIM) images to post-expansion confocal images, of the same sample, and calculating the distortion across the images. We imaged fixed U2OS cells expressing mitochondrial matrix-targeted GFP with SIM (see **Supplementary Methods** for details; **Supp. Fig. 2c**, without anti-GFP labeling; given that resolving the mitochondrial matrix vs. membrane requires 30 nm resolution[1,2], and a classical ExM (that expands ~4x) offers ~60-70 nm resolution[3,4], matrix-targeted GFP is indistinguishable from mitochondrial membrane in the context of the current experiment. We then performed mExM on, and imaged, the very same cells with a confocal microscope. Comparing pre-expansion SIM images of mitochondrial matrix-targeted GFP to post-expansion images of either GFP (with anti-GFP labeling for boosting GFP signals), or pGk5b, we observed the same low distortion (a few percent, over ~10 μm) as was found for previous ExM protocols (see **Supplementary Methods** for details; **Supp. Fig. 2f** and **Supp. Fig. 2i**). By comparing the distance between two landmarks in pre- vs. post-expansion images (**Supp. Fig. 2j** and **Supp. Fig. 2k**) of the same sample, the expansion factor could be calculated; we obtained an expansion factor (~4.4, **Supp. Fig. 2l**) similar to what was previously reported[3,4].

**Supplementary Note 3**

We expressed mitochondrial matrix-targeted GFP or endoplasmic reticulum (ER) membrane-targeted GFP in HEK293 cells via BacMam virus. We performed mExM on the cells (S**upp. Fig. 3a-b;** see **Supplementary Methods** for details), and then imaged the expanded cells, so that we could quantify the fraction of mitochondrial matrix- and ER membrane-targeted fluorescent protein signal that also exhibited pGk5b signal (see **Supplementary Methods** for details; in summary, a pixel was considered pGk5b-positive if it was brighter than one standard deviation below the pGk5b mean that was measured across the whole images). As a result, we observed that >99% of the mitochondrial matrix-targeted and ER membrane-targeted GFP signals also exhibited pGk5b signals (n=3 separate cells from 1 culture; **Supp. Fig. 3d**).

**Supplementary Note 4**

To enable post-expansion antibody staining, we adopted a commonly used ExM softening protocol[5] (i.e., SDS solution at a high temperature) that can reveal previously unseen structures by preserving protein epitopes through the expansion process[5]. This protocol builds upon post-expansion protein-retention ExM (proExM) protocols[4], as well as tissue proteomics protocols for formalin-fixed paraffin-embedded (FFPE) tissues[6,7]. In brief, we heat the sample for half an hour at 100 ºC and for 2 hours at 80 ºC, in a "fixation reversal" (FR) buffer[8] containing 0.5% PEG20000, 100mM DTT, 4% SDS, in 100mM Tris pH8 (see **Supp. Table 2** for details).

**Supplementary Note 5**

Using the softening solution (see **Supplementary Note 4**, **Supp. Table 2**), we performed mExM with antibody staining (see **Supplementary Methods** for details), using antibodies against organelle-specific membrane-localized proteins, TOM20 for mitochondria (**Supp. Fig. 5a**), and Nup98 for the nuclear pore complex (**Supp. Fig. 5b**). We also labeled myelin using an antibody against myelin basic protein (MBP) (**Supp. Fig. 5c**). We then quantified the fraction of signals from antibodies against membrane-localized proteins that also contained pGk5b signals, as we did for membrane-targeted GFP in cultured cells (see **Supplementary Note 3**). Since different antibodies may not react to the same sites on the same target protein[5], we used multiple antibodies against the same protein for TOM20 (two separate vendors) and MBP (three separate vendors) to further validate our technology.

**Supplementary Note 6**

The diameter of axons is known to be diverse across brain regions[9]. However, our finding aligns with measured axon diameters from EM images of the same brain regions (i.e., cortex and dentate gyrus)[9–12].

### Supplementary Figures

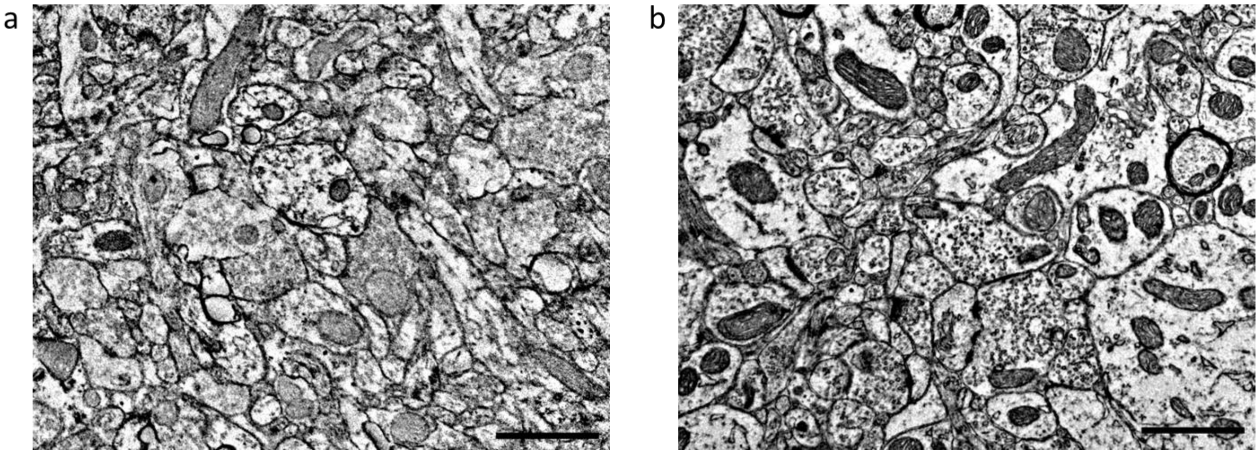

Supplementary Figure 1

Electron microscopy imaging of membrane label (pGk5a)-stained mouse brain slices (hippocampus region). In brief (see **Supplementary Methods** for details), 100 μM of palmitoylated glycine pentalysine peptide, equipped with an azide group (instead of biotin; termed pGk5a), without osmium counterstain (**a**), or no membrane probe but with osmium tetroxide counterstain (**b**), was applied to 100-μm thick tissue slices. Mouse brain tissue was preserved in 4% PFA and 0.1% glutaraldehyde at 4 ºC, and labeled with pGk5a for >16 hours at 4 ºC. The tissue was post-fixed in 2% PFA and 2% glutaraldehyde and labeled with 1.8nm undecagold gold nanoparticles, conjugated to dibenzocyclooctyne to attach to the azide handle on pGk5a. The tissue was counter-labeled with uranyl acetate, embedded in resin, sliced, and imaged on a TEM scope. Since the common practice of uranyl acetate (UA) staining without osmium does not clearly visualize the membrane (Fig. 3 from ref[13], Fig 4. from ref[14]), we reasoned that UA without osmium reacts to proteins as well as the amino groups (i.e., lysines) of pGk5a. We thus decided to use a common UA staining protocol (1% UA for 1 hour at room temperature) to enhance pGk5a signals on top of signals from gold nanoparticles. As a control, the tissue underwent the same protocol as described above, but without membrane probe incubation and with osmium. When labeling membranes with the membrane probe in the absence of the osmium counterstain, the stain appears to label the membranes as with osmium, but with slightly lower contrast. Scale bars: 1 μm.

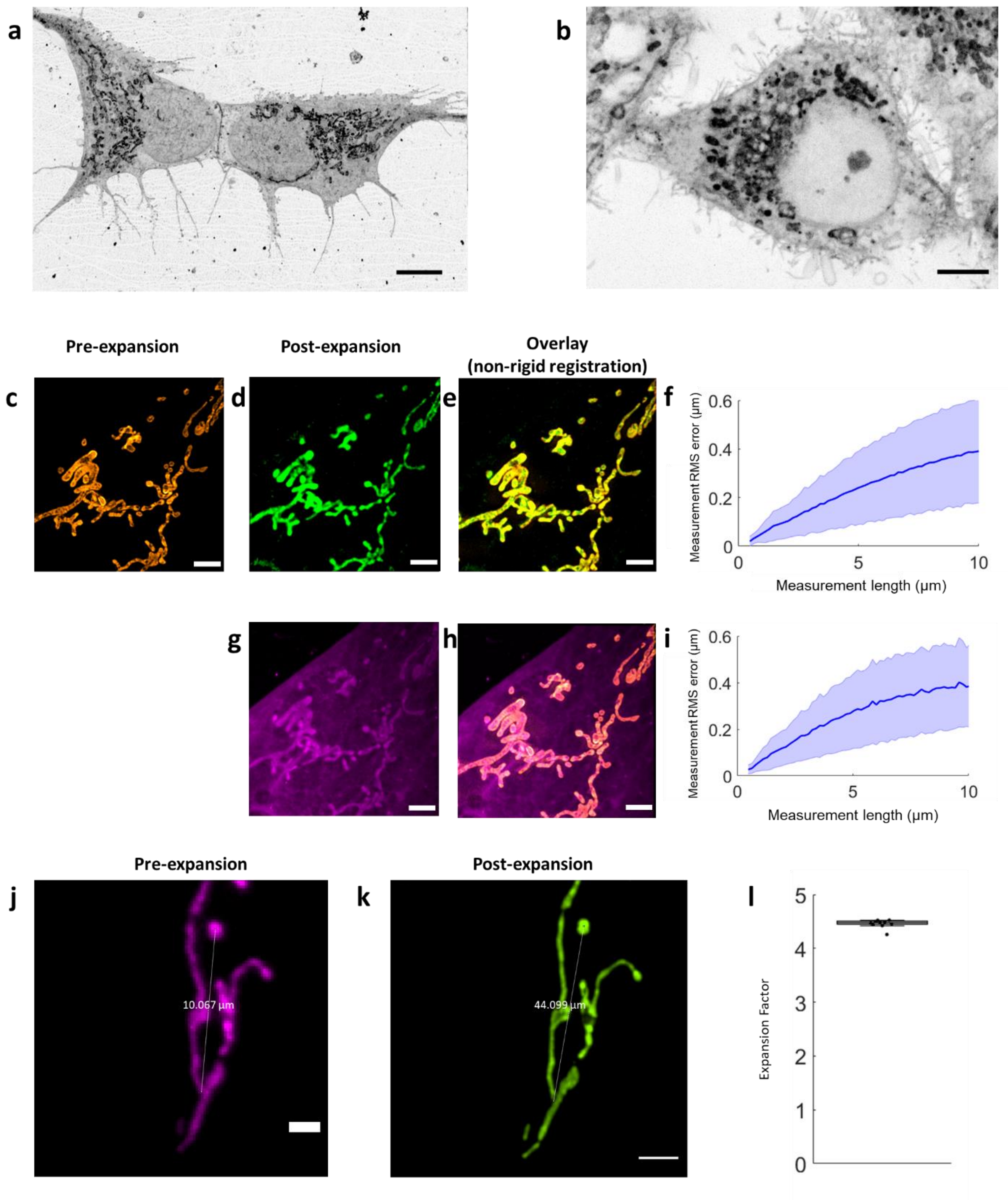

Supplementary Figure 2

(**a**) Representative (n=12 cells from two cultures) single z-plane confocal image of expanded HEK293 cell that underwent the mExM protocol (see **Supplementary Methods**), showing pGk5b staining of the membrane. Image visualized in inverted gray color (dark signals on light background). (**b**) Representative (n=12 cells from two cultures) single z-plane confocal image of expanded HeLa cell that underwent the mExM protocol, showing pGk5b staining of the membrane. Image visualized in inverted gray color. (**c**) Representative (n=3 cells from one culture) single z-plane structured illumination microscopy (SIM) image of a pre-expanded U2OS cell expressing mitochondrial matrix-targeted green fluorescent protein (GFP). Image visualized in orange color. (**d**) Single z-plane confocal image of the same U2OS cell as in (**c**), after undergoing the mExM protocol, showing expression of mitochondrial matrix-targeted GFP in the same field of view as shown in (**c**). Image visualized in green color. (**e**) non-rigidly registered and overlaid pre-expansion SIM image of the U2OS cell expressing mitochondrial matrix-targeted GFP in (**c**), and post-expansion confocal image of the same fixed U2OS cell after mExM processing, showing the mitochondrial matrix-targeted GFP channel in (**d**). (**f**) Root mean square (RMS) length measurement error as a function of measurement length, comparing pre-expansion SIM images of fixed U2OS cell expressing mitochondrial matrix-targeted GFP and post-expansion confocal images of the same cells after mExM processing, showing mitochondrial matrix-targeted GFP (blue line, mean; shaded area, standard deviation; n=3 cells from one culture). (**g**) Single z-plane confocal image of the same mExM-expanded fixed U2OS cell as in (**d**), showing pGk5b staining of the membrane in the same field of view as shown in (**c**). Image visualized in magenta color. (**h**) Non-rigidly registered and overlaid pre-expansion SIM image of the U2OS cell expressing mitochondrial matrix-targeted GFP in (**c**) and post-expansion confocal image of the same U2OS cell in (**e**) after mExM processing, showing pGk5b staining. (**i**) Root mean square (RMS) length measurement error as a function of measurement length, comparing pre-expansion SIM images of fixed U2OS cells expressing mitochondrial matrix-targeted GFP, and post-expansion confocal images of the same cell after mExM processing, showing pGk5b staining (blue line, mean; shaded area, standard deviation; n=3 cells from one culture). (**j-k**) Expansion factor analysis on HEK293 cells, that underwent mExM, after expressing mitochondrial matrix-targeted GFP. We randomly choose two landmark points in pre-expansion images and found the corresponding landmarks in expanded sample images. We then calculated the distance between the points, in both pre- and post-expansion images, and calculated the ratio to obtain the expansion factor. (**j**) Representative (out of 10 cells from two cultures) single z-plane confocal image of pre-expanded HEK293 cell. (**k**) As in (**j**), but post-expansion, for the same field of view shown in (**j**). (**l**) Boxplot showing measured expansion factor (n =10 cells from two cultures; black points, individual measured expansion factor, median, middle line; 1$^{st}$ quartile, lower box boundary; 3$^{rd}$ quartile, upper box boundary; error bars are the 95% confidence interval). Scale bars: (**a**-**b**) 5 μm, (**c**,**d**,**e**,**g**,**h,j**) 2 μm in biological units, (**k**) 10μm in post-expansion units.

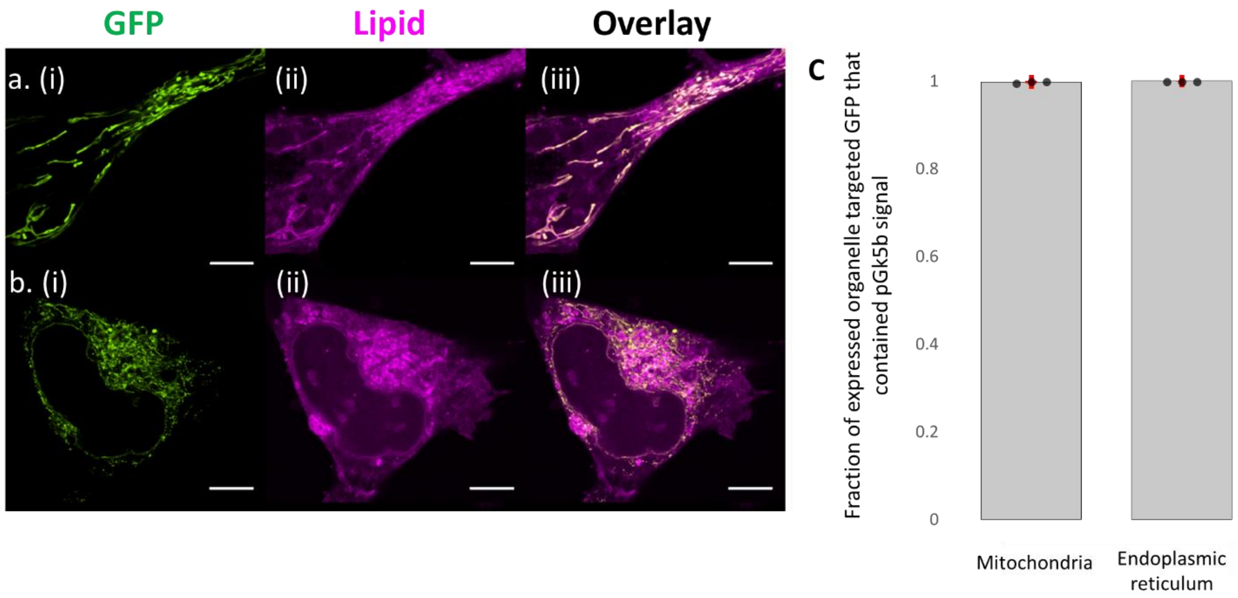

Supplementary Figure 3

(**a.i**) Representative (n=4 cells from one culture) single z-plane confocal image of expanded HEK293 cell expressing mitochondrial matrix-targeted GFP, after mExM processing. (**a.ii**) Single z-plane confocal image of the same expanded HEK293 cell as in (**a.i**), showing pGk5b staining in the same field of view as in (**a.i**). (**a.iii**) Overlay of (**a.i**) and (**a.ii**). (**b.**) As in (**a.**), but for a HEK293 cell expressing ER membrane-targeted GFP (n=3 cells from one culture). (**c**) Fraction of the pixels containing mitochondrial matrix-targeted GFP signal (left, n=3 cells from one culture; black points, individual data points, bar, mean, red error bars, standard deviation, used throughout unless otherwise noted) or ER membrane-targeted GFP signal (right, n=3 cells from one culture) that also exhibited pGk5b signal, in mExM-processed HEK293 cells (see **Supplementary Methods**). Scale bars are provided in biological units (i.e., physical size divided by expansion factor): (**a**-**b**) 5 µm.

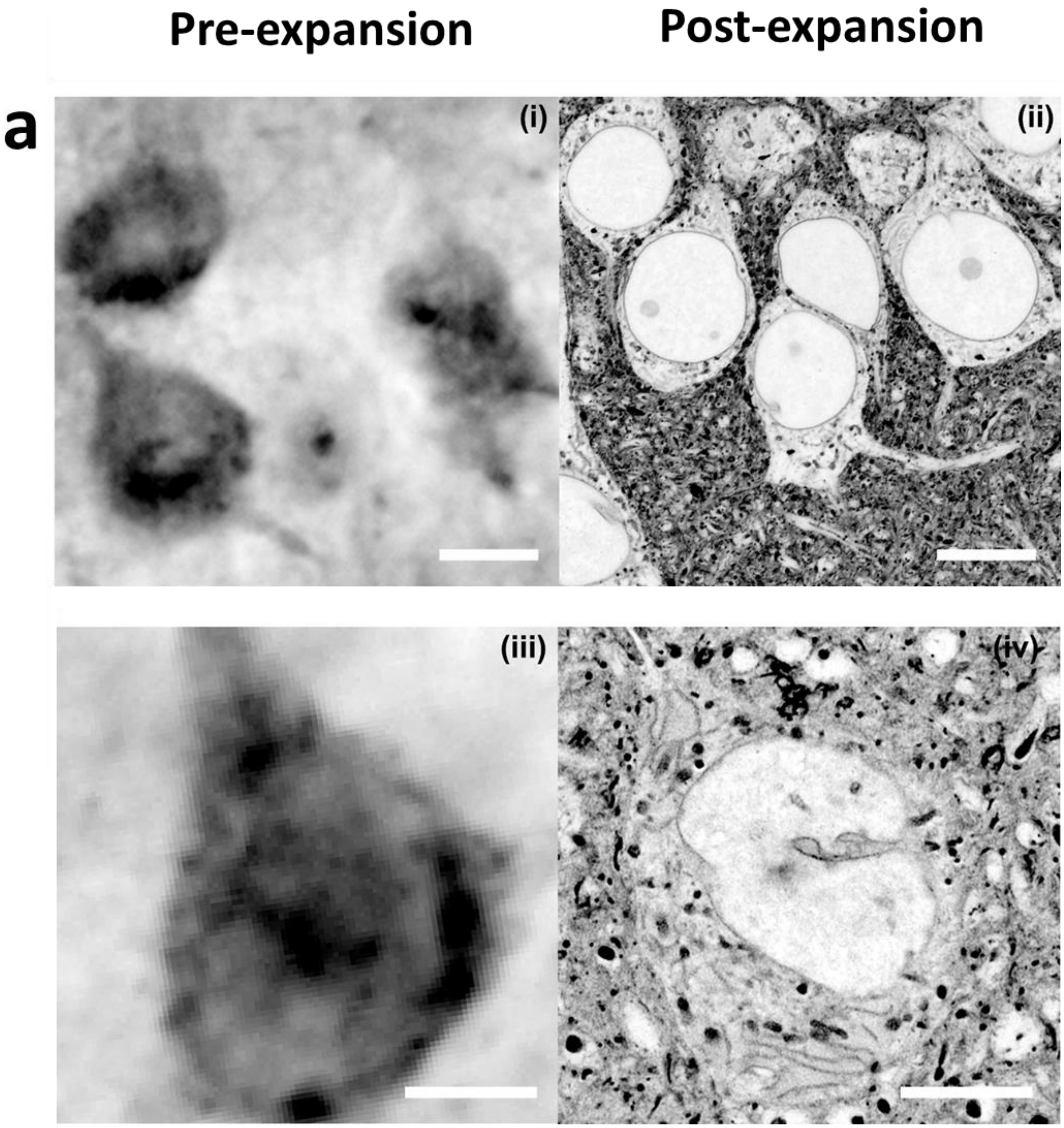

Supplementary Figure 4

(**a.i**) Representative (n=5 fixed brain slices from two mice) single z-plane confocal image of pre-expanded mouse brain tissue (cortex) showing pGk5b staining. Images visualized in inverted gray color throughout this figure (dark signals on light background). (**a.ii**) Representative (n=5 fixed brain slices from two mice) single z-plane confocal image of expanded mouse brain tissue after mExM processing, showing pGk5b staining. (**a.iii**) As in (**a.i**) but focused on one cell body. (**a.iiii**) As in (**a.ii**) but focused on one cell body. Scale bars are provided in biological units (i.e., physical size divided by expansion factor): (**a.i**) 10 μm, (**a.ii**) 10 μm, (**a. iii**) 5 μm, (**a. iv**) 5 μm.

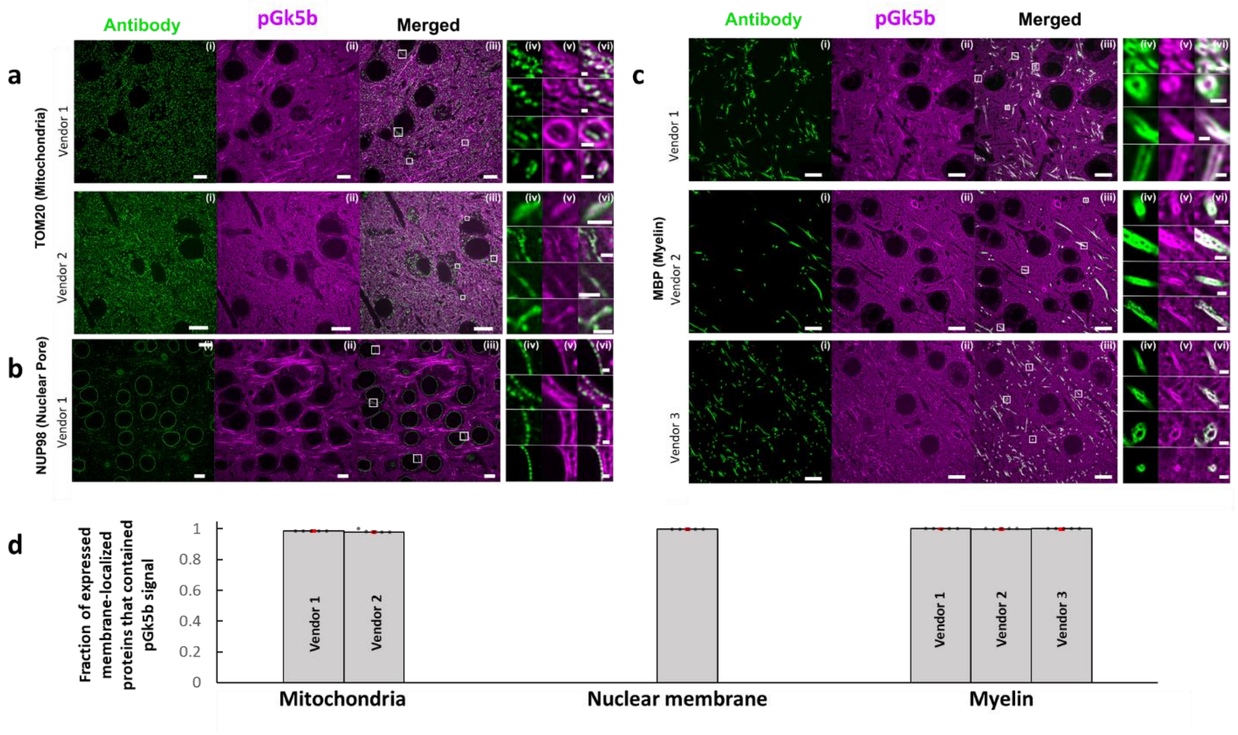

Supplementary Figure 5

(**a.i**) Representative (n=5 cortex or hippocampus regions from the same mouse brain) single z-plane confocal image of expanded mouse brain tissue after mExM processing with immunostaining, showing immunolabeling against the mitochondrial membrane protein Tom20, using antibodies from two separate vendors (see **Supp Table 2** for details). (**a.ii**) Single z-plane confocal image of the specimen of (**a.i**), showing pGk5b staining of the same field of view as in (**a.i**). (**a.iii**) Overlay of (**a.i**) and (**a.ii**). (**a.iv**) Magnified views of white boxed regions in (**a.iii**) showing TOM20 signal (green). (**a.vi**) As in (**a.iv)**, but showing pGk5b signal (magenta). (**a.vii**) Overlay of (**iv**) and (**v**). (**b.**) As in (**a.**), but for the nuclear pore protein NUP98. (**c.**) As in (**a.**), but for myelin basic protein (MBP), using antibodies from three separate vendors (see **Supp Table 2** for details). (**d.**) Fraction of the pixels containing each membrane protein signal, that also contained pGk5b signal, in mExM-processed mouse brain tissue, for each of the antibodies used above (see **Supplementary Methods**; for each bar, n=5 cortex or hippocampus regions from the same mouse brain). Scale bars: (**i-iii**) 10μm, (**iv-vi**) 1μm.

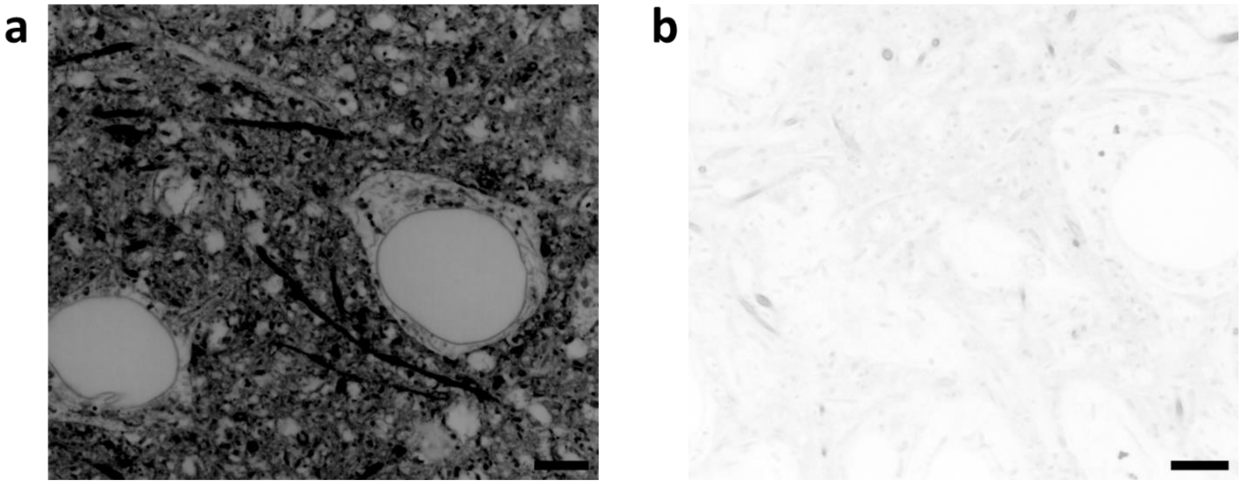

Supplementary Figure 6

We tested two versions of the membrane labeling probe with ExM: **a**) palmitoyl-glycine-(D-lysine)$_5$-biotin (pGk5b) and **b**) farnesyl-glycine-(D-lysine)$_5$-biotin (i.e., replacing palmitoyl in pGk5b with farnesyl). (**a**) Representative (n=5 fixed brain slices from two mice) single z-plane confocal image of expanded mouse brain tissue (hippocampus) after mExM processing, showing pGk5b (i.e., the palmitoylated form of the membrane labeling probe) staining of the membrane. The image is visualized in inverted gray color. (**b**) as in (**a**), but with a farnesylated form of the membrane labeling probe, showing the farnesylated form of the probe staining of the membrane. The image is visualized in inverted gray color. Images (**a**-**b**) are visualized with the same brightness and contrast with ImageJ software to highlight the difference between the two images. Scale bars are provided in biological units (i.e., physical size divided by expansion factor): (**a, b**) 5 μm.

a

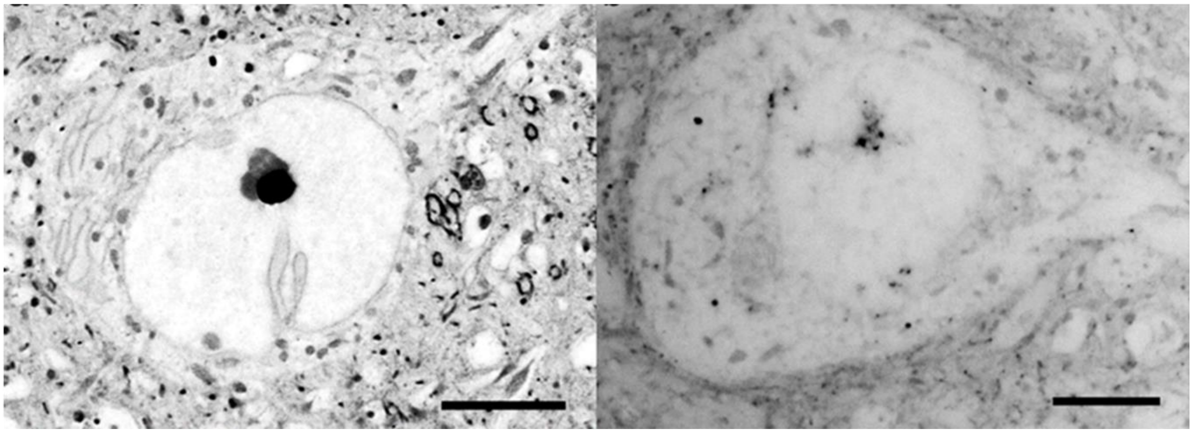

Supplementary Figure 7

(**a**) Effect of the glycine linker attached to the palmitoyl group on the efficacy of membrane labeling in fixed brain tissue. Image visualized in inverted gray color. We tested two versions of the palmitoylated 5 lysine biotin membrane probe: left) one containing a glycine linker attached to the palmitoyl group enabling flexibility of the lipid relative to the peptide carrier, and right: one that does not contain a glycine but in which the lipid is directly attached to the lysine backbone. In the case of the glycine linker, the level of detail we achieve in labeling membranes is superior to that achieved without the glycine linker. Scale bars are provided in biological units (i.e., physical size divided by expansion factor): (**a-b**) 5 μm.

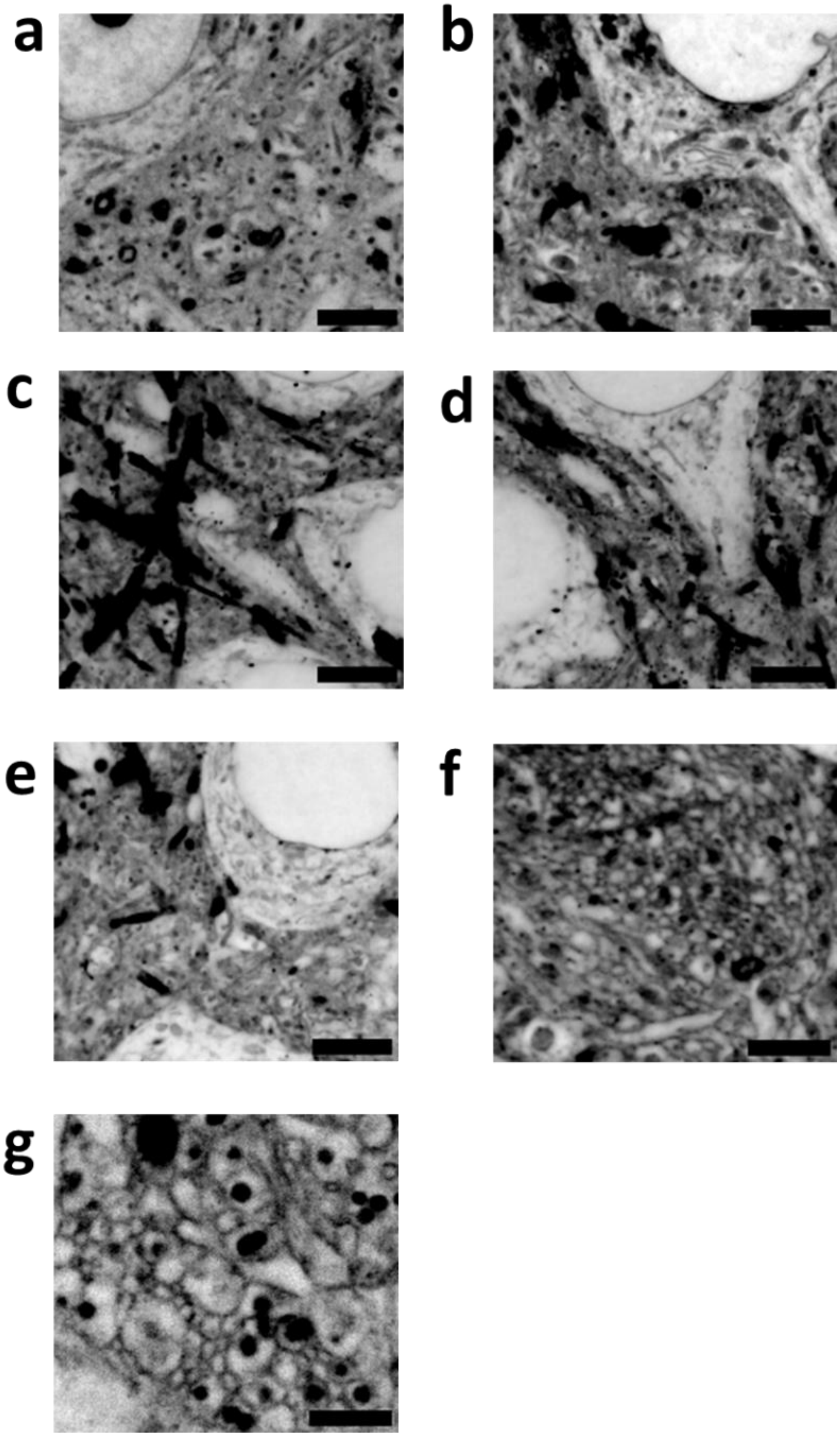

#### Supplementary Figure 8

We tested varying numbers of lysines (i.e., 3, 5, 7, 9, 11, 13, and 19 lysines) in the backbone of the membrane labeling probe while holding other moieties (palmitoyl tail, glycine, and biotin) constant. (**a**) Representative (n=3 fixed brain slices from two mice) single z-plane confocal image of expanded mouse brain tissue (hippocampus) after mExM processing, but with a membrane labeling probe containing 3 lysines, showing the probe staining of the membrane. The image is visualized in inverted gray color. (**b**) - (**g**) As in (**a**) but with membrane labeling probes containing 5, 7, 9, 11, 13, and 19 lysines, respectively. Scale bars are provided in biological units (i.e., physical size divided by expansion factor): (**a-g**) 2 μm.

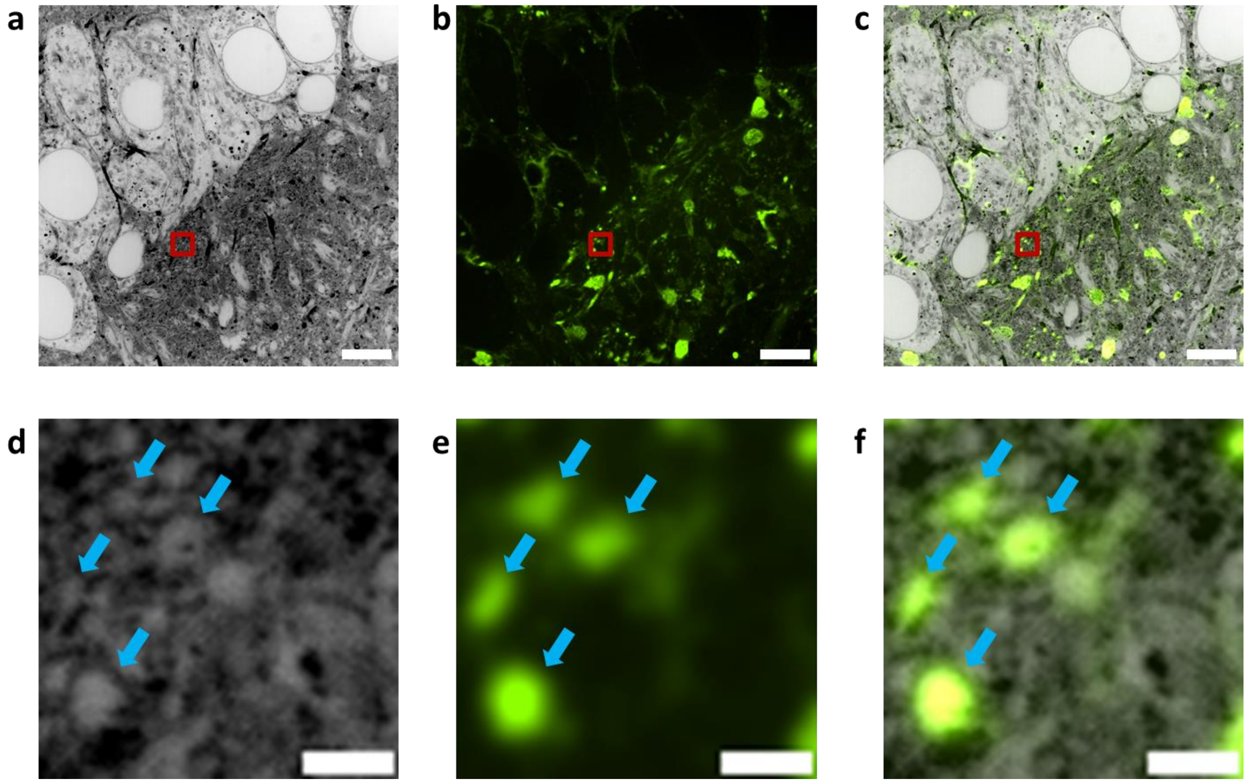

Supplementary Figure 9

(**a**) Representative (n=5 fixed brain slices from two mice) single z-plane confocal image of expanded Thy1-YFP mouse brain tissue (hippocampus) after mExM processing using pGk13b and stained with an anti-GFP antibody to boost the YFP signal, showing the pGk13b staining of the membrane. The image is visualized in inverted gray color. (**b**) Single z-plane confocal image of the specimen of (**a**) showing anti-GFP staining of the same field of view as in (**a**). (**c**) Overlay of (**a**) and (**b**). (**d**) Magnified views of red boxed region in (**a**). (**e**) Magnified views of red boxed region in (**b**). (**f**) Magnified views of red boxed region in (**c**). Scale bars are provided in biological units (i.e., physical size divided by expansion factor): (**a-c**) 5μm, (**d-f**) 0.5μm.

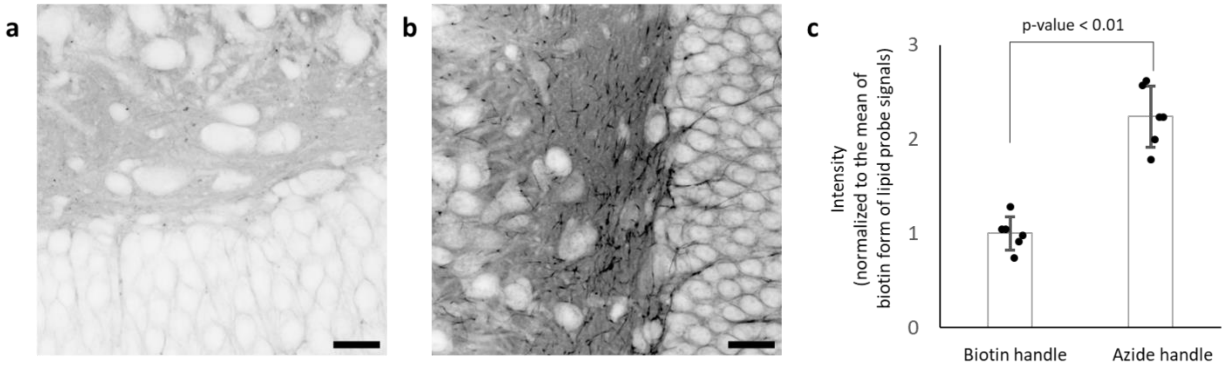

#### Supplementary Figure 10

The performance of a biotin handle (pGk13b) vs. azide handle (pGk13a) with fixed mouse brain tissues in the context of ExM. Mouse brain tissues were fixed with ice-cold 4% PFA. We applied pGk13b or pGk13a overnight, we then followed the standard ExM protocol[4]. In short, the tissues were processed with AcX, and the ExM gel was formed. After the tissue softening with proteinase K, the probe was fluorescently labeled with fluorescent streptavidin (i.e., cy3-streptavidin, >1 fluorophore per streptavidin), expanded, and imaged (**a**) or with fluorescent DBCO (i.e., cy3-DBCO, 1 fluorophore per DBCO), expanded and imaged (**b**). (**a**) Representative (n = 6 fixed brain slices from two mice) single z-plane confocal image of expanded mouse brain tissue (hippocampus), showing the pGk13b staining of the membrane. The image is visualized in inverted gray color. (**b**) as in (**a**) but with pGk13a, showing the pGk13a staining of the membrane. Images (**a**-**b**) are visualized with the same brightness and contrast with ImageJ software to highlight the difference between the two images. (**c**) The signal intensity of the pGk13b (left bar; n=6 fixed brain slices from two mice) and the pGk13a (right bar; n=6 fixed brain slices from two mice). Black points, individual measured average intensity of each image; bar, mean; error bars, standard deviation; p-value, unpaired two-sided t-test between signals from the pGk13b (left bar) and the pGk13a handle probe (right bar). Scale bars are provided in biological units (i.e., physical size divided by expansion factor): (**a, b**) 20 µm.

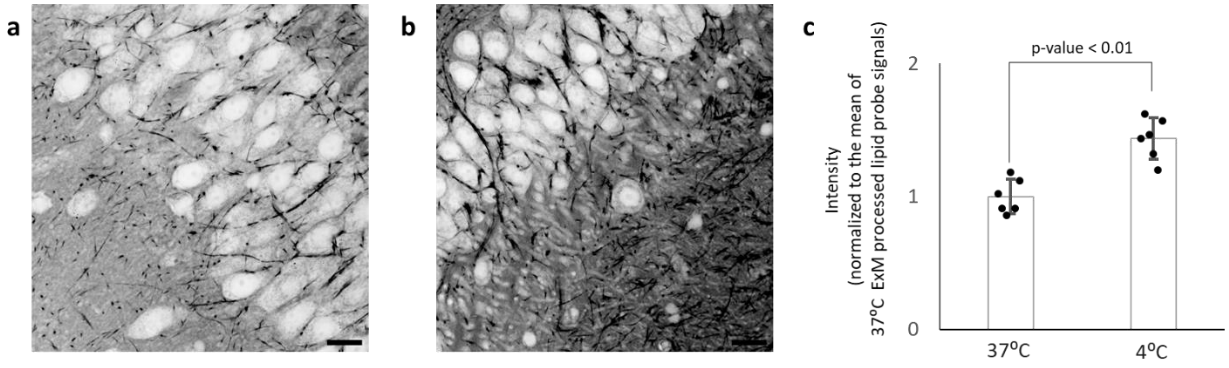

**Supplementary Figure 11**

pGk13a staining of the membrane of mouse brain tissue fixed with 4%PFA and 0.5% CaCl2 at 4 ºC and proceeded with the standard ExM protocol (37 ºC gelation)[4] vs. modified ExM protocol (i.e., 4 ºC gelation). Mouse brain tissues were fixed with ice-cold 4% PFA and 0.5% CaCl2 fixatives. We applied the pGk13a probe overnight at 4 ºC. We then performed the standard ExM protocol or modified ExM protocol (i.e., 4 ºC gelation instead of 37 ºC gelation). In short, the tissues were processed with AcX, and ExM gel was formed at 37 ºC (**a**) or 4 ºC (**b**). After the tissue softening with proteinase K, the pGk13a was fluorescently labeled with fluorescent DBCO (i.e., cy3-DBCO, 1 fluorophore per DBCO), expanded, and imaged (**a**-**b**). (**a**) Representative (n=6 fixed brain slices from two mice) single z-plane confocal image of expanded mouse brain (hippocampus) tissue showing pGk13a staining of the membrane after the standard ExM protocol. The image is visualized in inverted gray color. (**b**) as in (**a**) but with 4 ºC gelation. Images (**a**-**b**) are visualized with the same brightness and contrast with ImageJ software to highlight the difference between the two images. (**c**) The intensity of 37 ºC gelation (left bar; n=6 fixed brain slices from 2 mice) and 4 ºC gelation (right bar; n=6 fixed brain slices from two mice). Black points, individual measured average intensity of each image; bar, mean; error bars, standard deviation; p-value, unpaired two-sided t-test between signals from the 37 ºC gelation (left bar), and 4 ºC gelation (right bar). Scale bars are provided in biological units (i.e., physical size divided by expansion factor): (**a, b**) 20 μm.

a

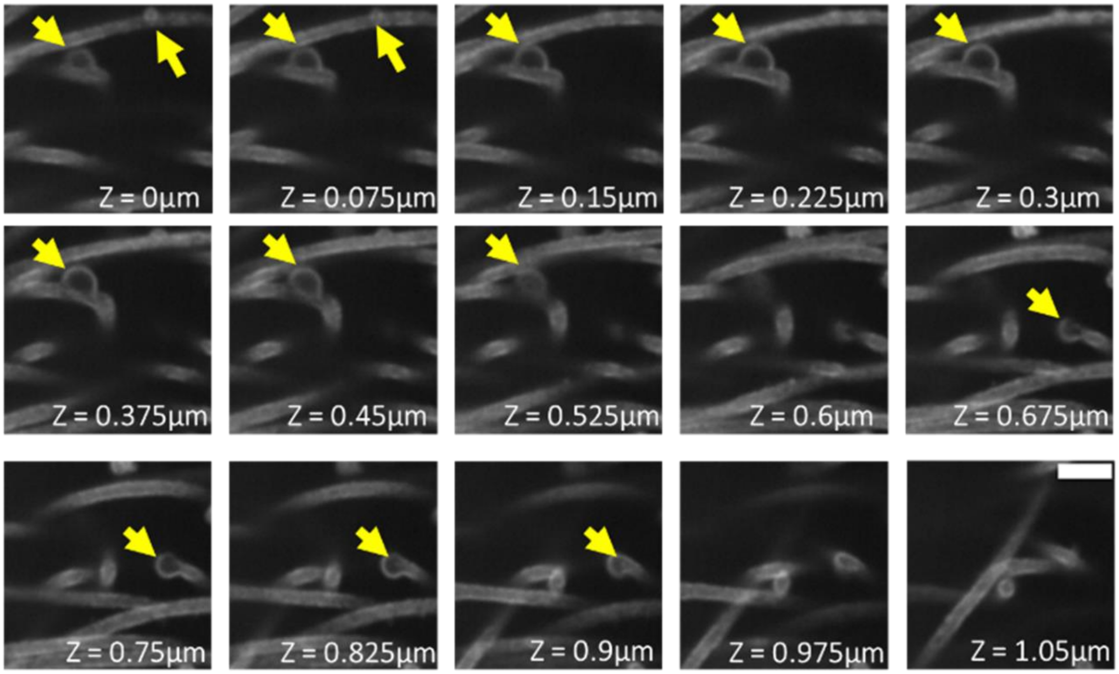

Supplementary Figure 12

(**a**) Fifteen serial sections from the 3D volume rendering in **Fig. 2s, right**. The yellow arrows indicate membrane vesicles. Scale bar in biological units (i.e., physical size divided by expansion factor): 1 μm.

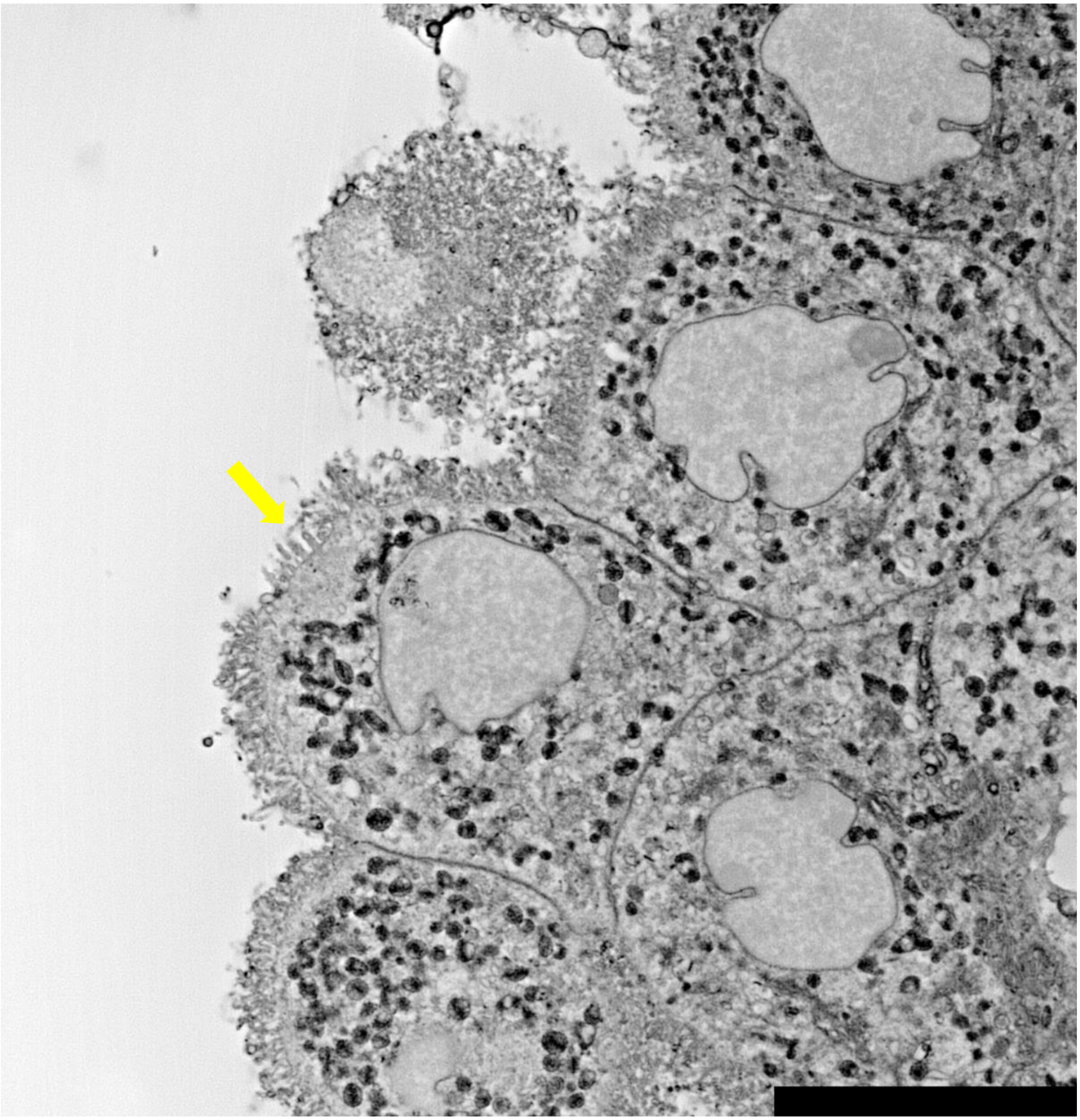

Supplementary Figure 13

Representative (n=5 fixed brain slices from two mice) single z-plane confocal image of expanded mouse brain tissue after umExM processing, showing the pGk13a staining of the membrane in the choroid plexus region. The image is visualized in inverted gray color. Example microvilli in the choroid plexus are pointed at with a yellow arrow. Scale bar in biological units (i.e., physical size divided by expansion factor): 10 μm.

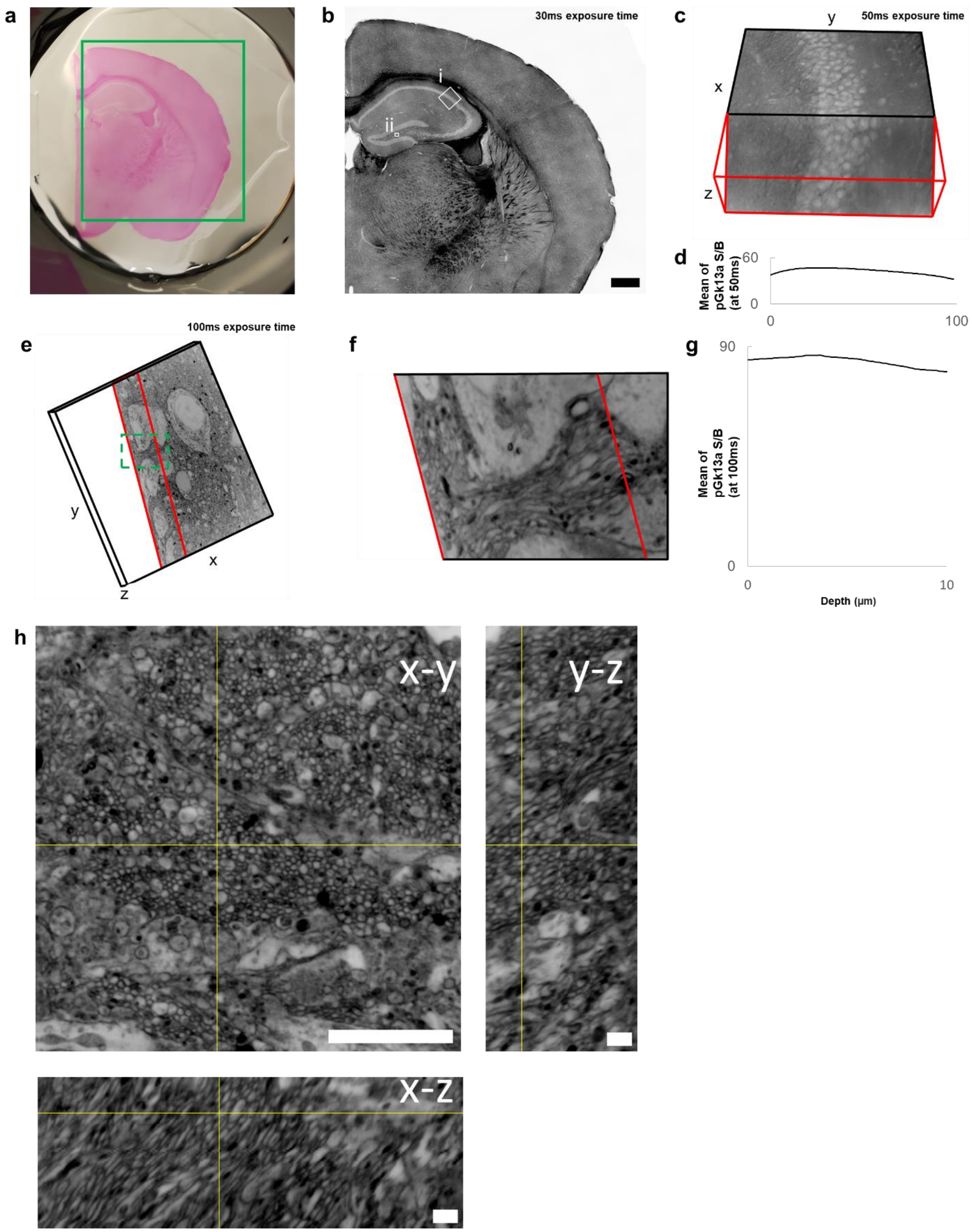

Supplementary Figure 14

(**a**) Photograph of a fixed 100µm thick adult mouse coronal slice that underwent the umExM protocol. (**b**) Single z-plane confocal image of green boxed region in (**a**). Images are taken with a

4x objective at 30ms laser exposure time, and they were stitched with shading correction function via default setting from Nikon Element software version 4.30. pGk13a staining of the membrane visualized in inverted gray color throughout this figure (dark signals on light background). We did not perform any image processing (e.g., denoising or deconvolution) other than stitching for images presented throughout this figure. (**c**) Volume rendering of the white box (i) in (**b**). Images were taken with a 4x objective at 50ms laser exposure time with a z step size of 0.375μm (in biological unit). Unless otherwise noted, clipping planes that are red colored indicate the portion that has been clipped out to expose the inside of the volume for 3D images presented throughout this figure. (**d**) Profile of mean pGk13a signal intensity of XY planes taken along the depth of the volume in (**c**). (**e**) Volume rendering of the white box (ii) in (**b**). Images were taken with a 60x objective at 100ms laser exposure time with a z step size of 0.125μm. (**f**) Magnified view of green boxed region in (**e**). (**g**) Profile of mean pGk13a signal intensity of XY planes taken along the depth of the volume in (**e**). (**h**) Cross-sectional images of dentate gyrus region in 100-μm thick mouse coronal slices that underwent the umExM protocol, showing pGk13a staining of the membrane. Images are taken with a 60x objective at 100ms laser exposure time with a z step size of 0.075μm. Yellow lines indicate the cross-sectional views in y-z and x-z images. Scale bars are provided in biological units (i.e., physical size divided by expansion factor) (**b**) 500μm, (**c**) 340μm (x); 340μm (y); and 100μm (z), (**e**) 62μm (x); 62μm (y); and 20μm (z) (**h**) 5μm (x-y); 1μm (y-z); 1μm (x-z).

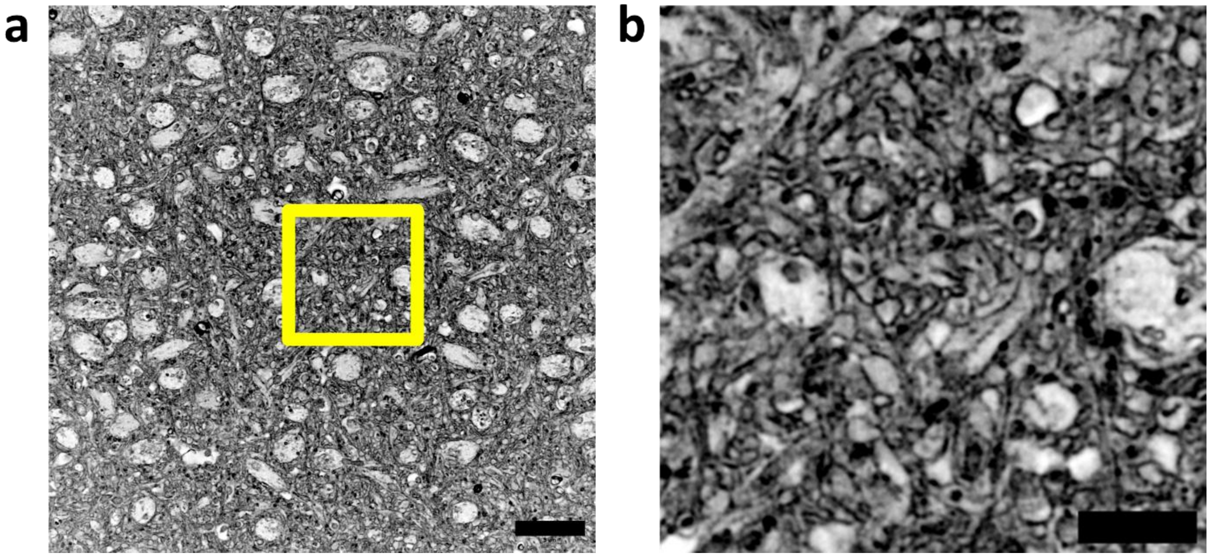

Supplementary Figure 15

(**a**) Representative (n=5 fixed brain slices from two mice) single z-plane confocal image of expanded mouse brain tissue after umExM processing, showing pGk13a staining of the membrane in hippocampal CA2. The image is visualized in inverted gray color. (**b**) Magnified view of yellow boxed region in (**a**). The image is visualized in inverted gray color. Scale bars are provided in biological units (i.e., physical size divided by expansion factor): (**a**) 10 μm, (**b**) 2 μm.

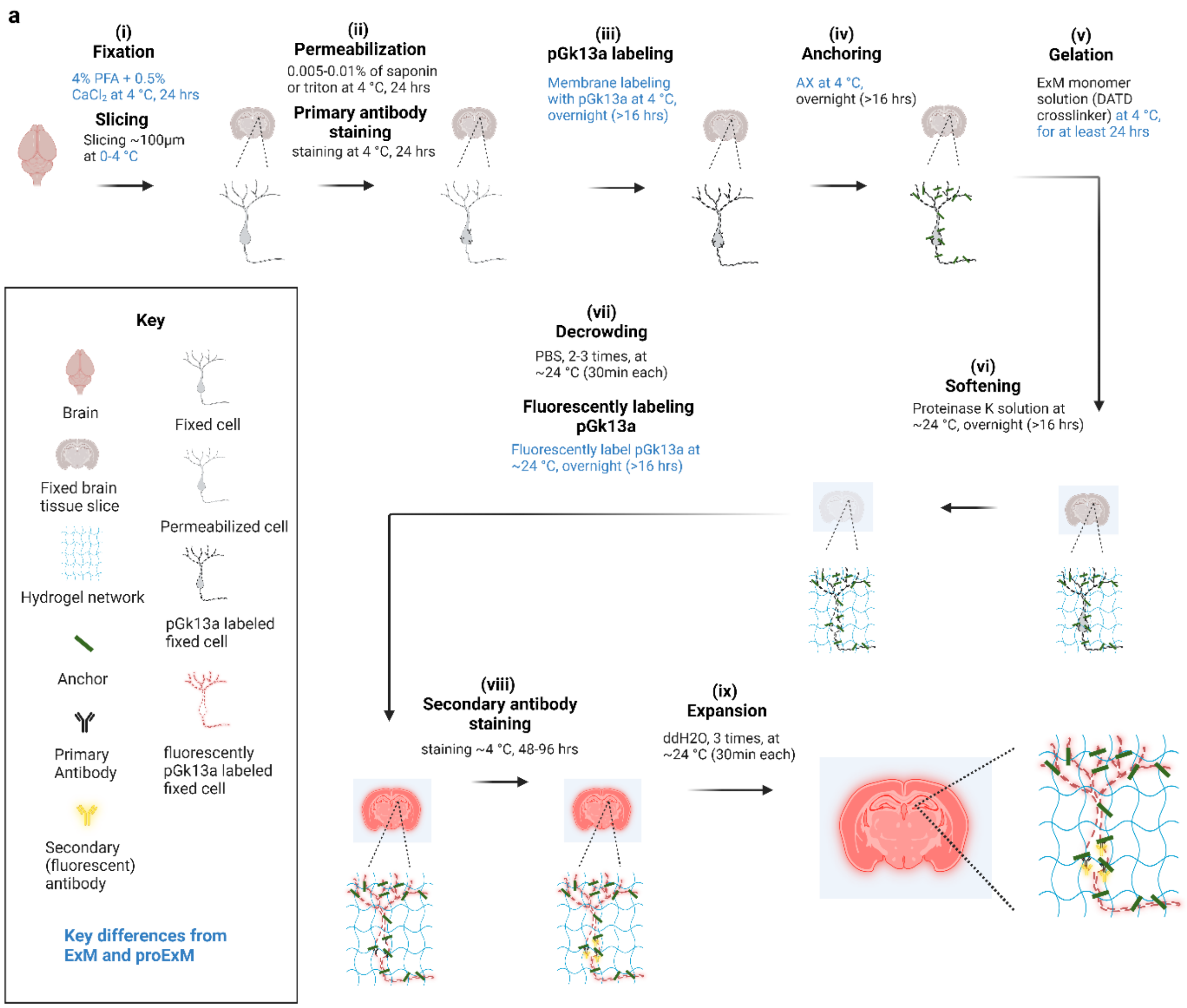

#### Supplementary Figure 16

(**a**) umExM with pre-expansion antibody staining workflow schematic. Blue-colored captions highlight the key differences from ExM[3] and proExM[4], whereas black captions highlight steps similar to those of earlier protocols. (**a.i**) A specimen is chemically fixed with 4% PFA + 0.5% $CaCl_2$ at 4 ºC for 24 hours. The brain is sliced on a vibratome at 100 μm thickness at 4 ºC. (**a.ii**) The specimen is treated with either 0.005-0.01% detergent (i.e., saponin or triton) at 4 ºC overnight (unless otherwise noted, overnight means >16 hours throughout the figure). Then specimen is incubated with a primary antibody. (**a.iii**) The specimen is treated with pGk13a at 4 ºC overnight. (**a.iv**) The specimen is treated with AX at 4 ºC overnight. (**a.v**) The specimen is embedded in an expandable hydrogel (made with DATD crosslinker[15]) at 4 ºC overnight. (**a.vi**) The sample (specimen-embedded hydrogel) is chemically softened with enzymatic cleavage of proteins (i.e., non-specific cleavage with proteinase K) at room temperature (~24 ºC), overnight. (**a.vii**) Then, the sample is treated with PBS to partially expand it. The pGk13a, that is anchored to the gel matrix, is fluorescently labeled via click-chemistry (i.e., DBCO-fluorophore) at room temperature, overnight. (**a.viii**) Then the sample is incubated with a secondary antibody at 4 ºC for 2-3 days. (**a.ix**) The sample is expanded with water at room temperature for 1.5 hours (exchanging water every 30 minutes).

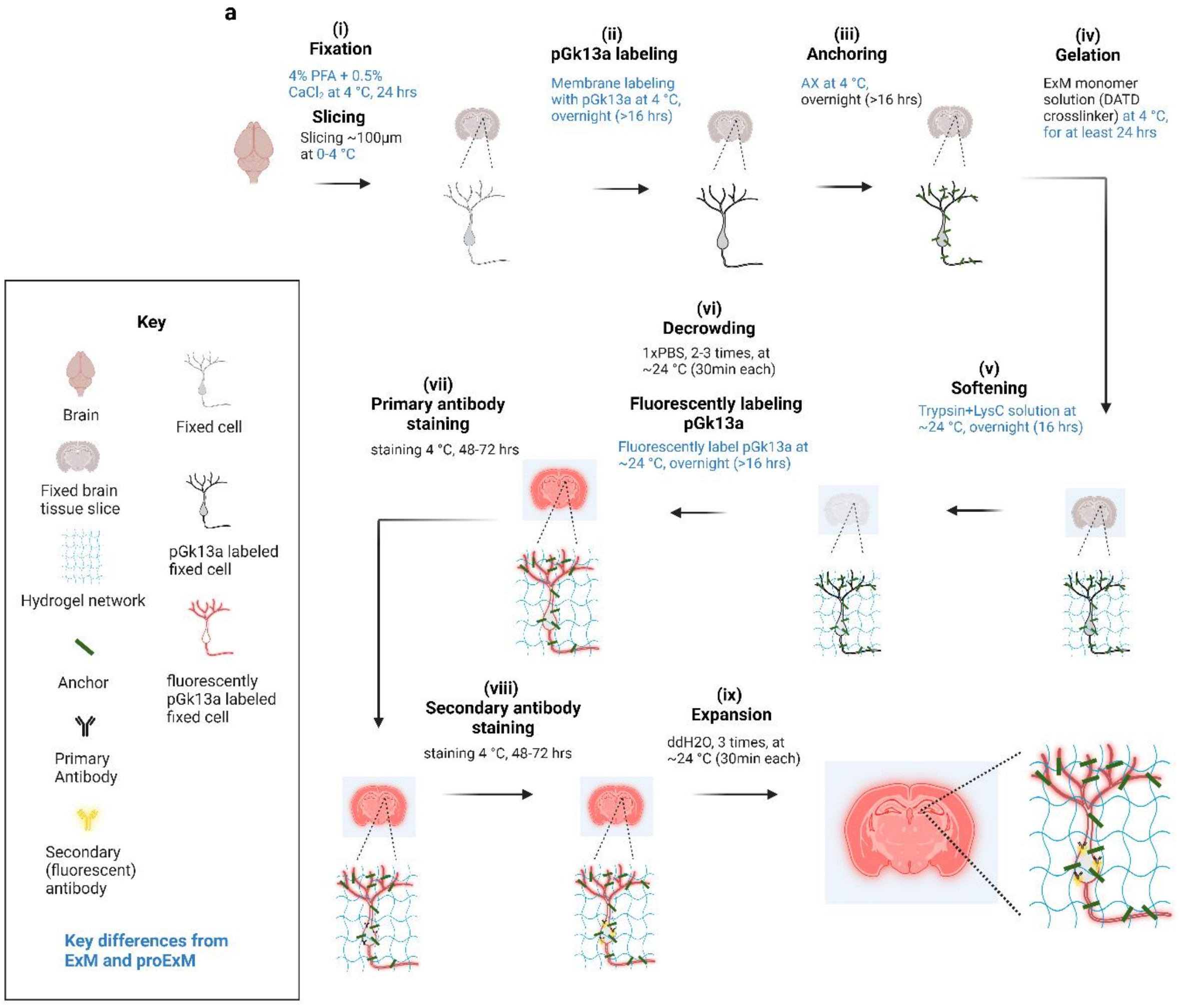

Supplementary Figure 17

(**a**) umExM with post-expansion antibody staining workflow schematic. Blue-colored captions highlight the key differences from ExM[3] and proExM[4], whereas black captions highlight steps similar to those of ExM and proExM. (**a.i**) A specimen is chemically fixed with 4% PFA + 0.5% $CaCl_2$ at 4 ºC for 24 hours. The brain is sliced on a vibratome at 100μm thickness at 4 ºC. (**a.ii**) The specimen is treated with pGk13a at 4 ºC overnight (unless otherwise noted, overnight means >16 hours throughout the figure). (**a.iii**) The specimen is treated with AX at 4 ºC overnight. (**a.iv**) The specimen is embedded in an expandable hydrogel (made with DATD crosslinker[15]) at 4 ºC overnight. (**a.v**) The sample (specimen-embedded hydrogel) is mechanically softened with enzymatic cleavage of proteins (i.e., specific cleavage with Trypsin and LysC) at room temperature (~24 ºC), overnight. (**a.vi**) Then, the sample is treated with PBS to partially expand it. The pGk13a, that is anchored to the gel matrix, is fluorescently labeled via click-chemistry (i.e., DBCO-fluorophore) at room temperature, overnight. (**a.vii**) Then the sample is incubated with a primary antibody at ~4 ºC, for 48-72 hours. (**a.vii**) Then the sample is incubated with a secondary antibody at ~4 ºC, for 48-48 hours. (**a.ix**) The sample is expanded with water at room temperature for 1.5 hours (exchanging water every 30 minutes).

**a.**

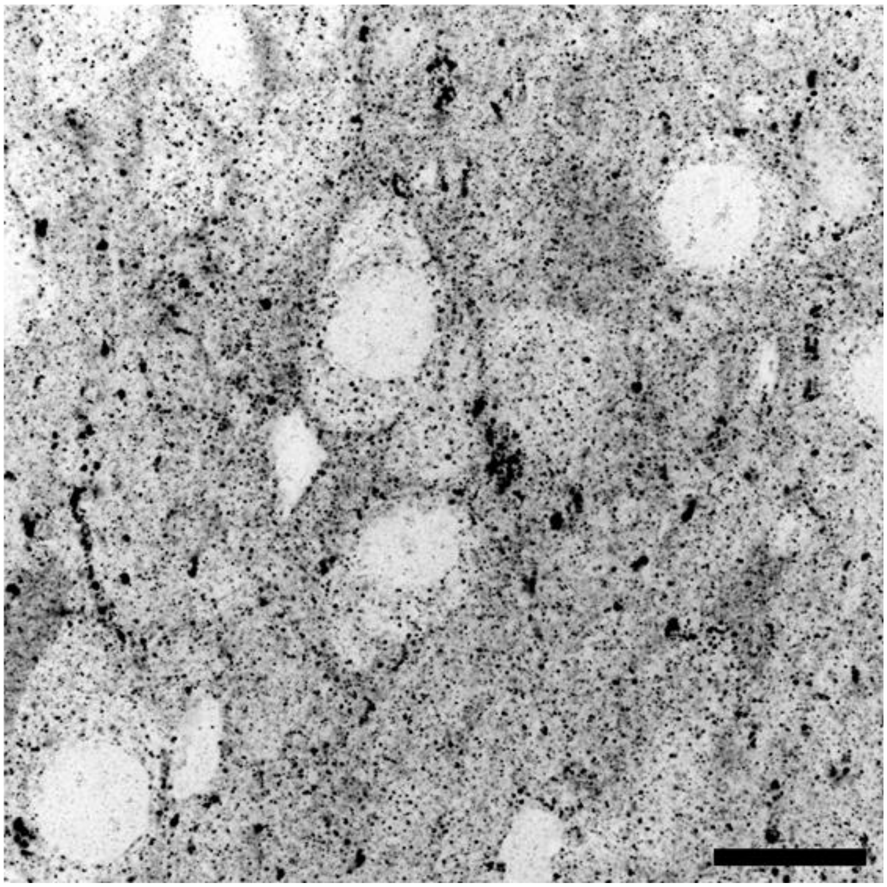

Supplementary Figure 18

(**a**) Representative (n=2 fixed brain slices from one mouse) single z-plane confocal image of expanded mouse brain tissue that underwent umExM protocol with the GMA anchor instead of AX anchor, showing pGk13a staining of the membrane in the hippocampus dentate gyrus. Scale bars are provided in biological units (i.e., physical size divided by expansion factor): (**a**) 10 µm

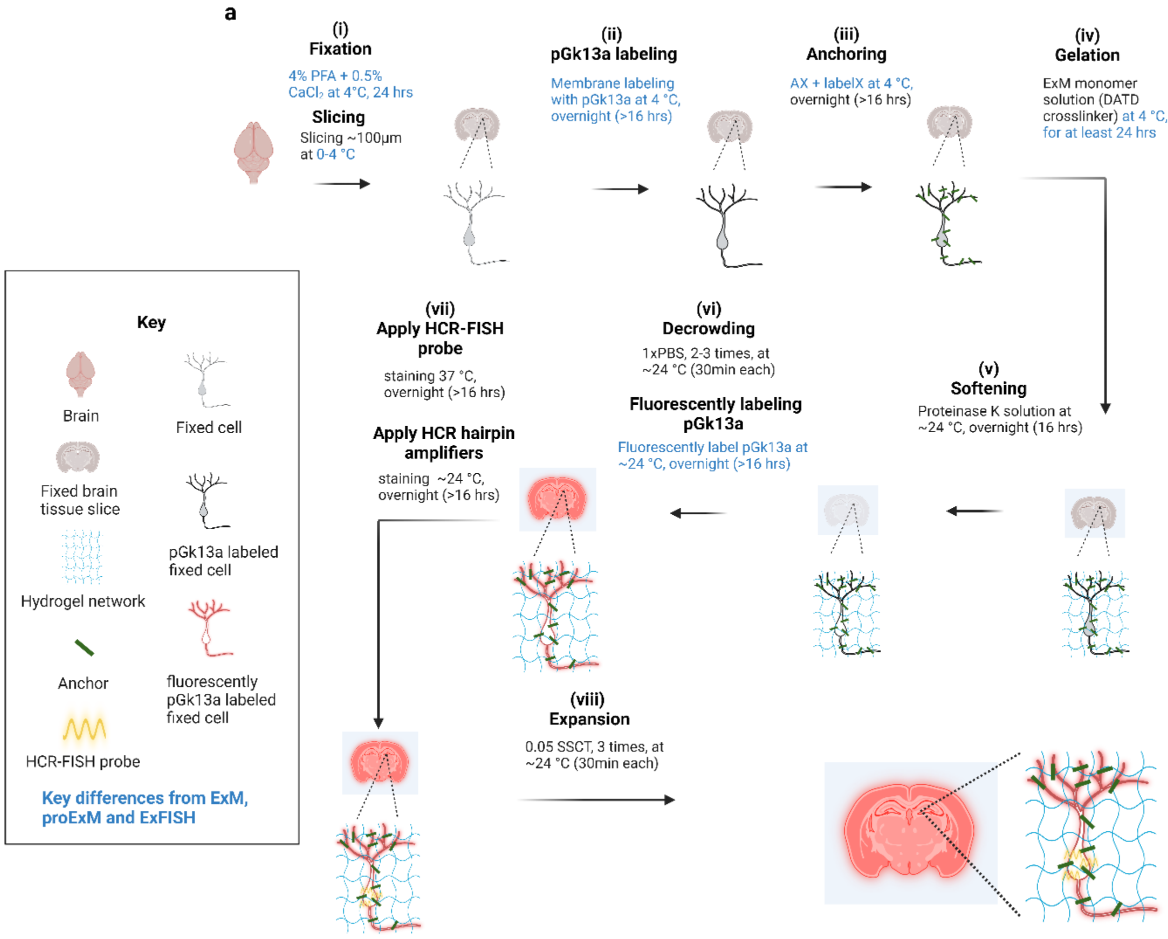

#### Supplementary Figure 19

(**a**) umExM with FISH workflow schematic. Blue-colored captions highlight the key differences from ExM[3], proExM[4] and ExFISH[16], whereas black captions highlight steps similar to those of ExM, proExM and ExFISH. (**a.i**) A specimen is chemically fixed with 4% paraformaldehyde (PFA) + 0.5% calcium chloride ($CaCl_2$) at 4 ºC for 24 hours. The brain is sliced on a vibratome at 100 µm thickness at 4 ºC. (**a.ii**) The specimen is treated with the pGk13a at 4 ºC overnight (unless otherwise noted, overnight means >16 hours throughout the figure). (**a.iii**) The specimen is treated with acrylic acid N-hydroxysuccinimide ester (AX) at 4 ºC overnight. (**a.iv**) The specimen is embedded in an expandable hydrogel (made with DATD crosslinker[15]) at 4 ºC for at least 24 hours. (**a.v**) The specimen is mechanically softened with enzymatic cleavage of proteins (i.e., specific cleavage with Proteinase k) at room temperature (~24 ºC), overnight. (**a.vi**) Then, the specimen-embedded hydrogel is treated with PBS to partially expand it. The pGk13a, that is anchored to the gel matrix, is fluorescently labeled via click-chemistry (i.e., DBCO-fluorophore) at room temperature, overnight. (**a.vii**) Then, specimen-embedded hydrogel is incubated with HCR-FISH probe at 37 ºC, overnight. (**a.vii**) Then, specimen-embedded hydrogel is incubated with fluorescently labeled HCR-hairpin amplifiers at ~24 ºC, overnight. (**a.viii**) The specimen-embedded hydrogel is expanded with 0.05x SSCT at room temperature for 1.5 hours (exchanging water every 30 minutes).

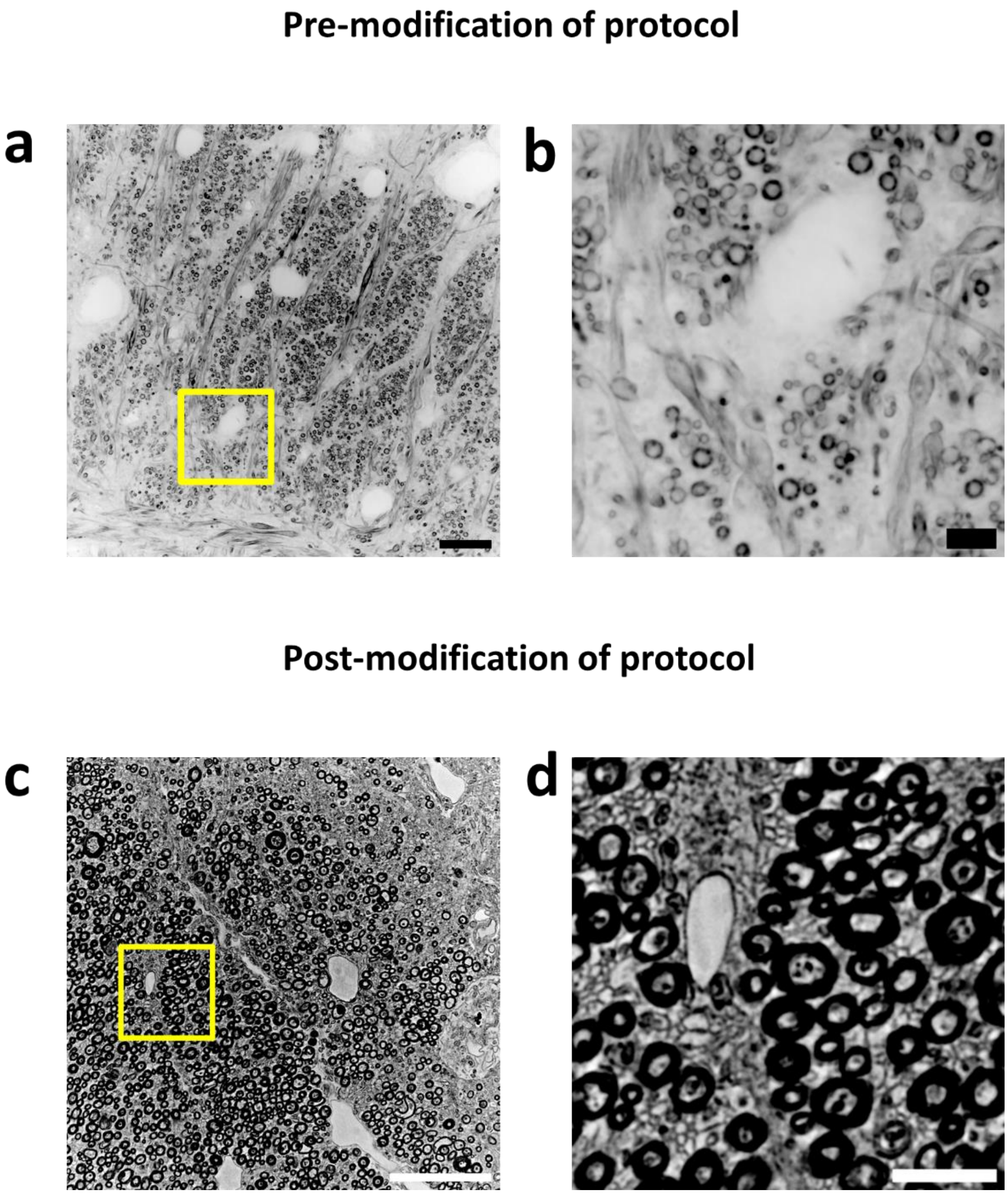

Supplementary Figure 20

(**a**) Representative (n=5 brain tissue sections from two mice) single z-plane confocal image of expanded mouse brain tissue after umExM protocol processing, showing pGk13a staining of the membrane in the corpus callosum region. The image is visualized in inverted gray color. (**b**) Magnified view of yellow boxed region in (**a**). Only a subset of axons can be identified in the images. (**c**) Representative (n=5 brain tissue sections from 2 mice) single z-plane confocal image of expanded mouse brain tissue after modified umExM protocol processing, showing pGk13a staining of the membrane in the corpus callosum region. The image is visualized in inverted gray color. (**d**) Magnified view of yellow boxed region in (**c**). The modification of the protocol drastically improved the visualization of axons in the corpus callosum. Scale bars are provided in biological units (i.e., physical size divided by expansion factor): (**a,c**) 10 μm, (**b**,**d**) 2 μm.

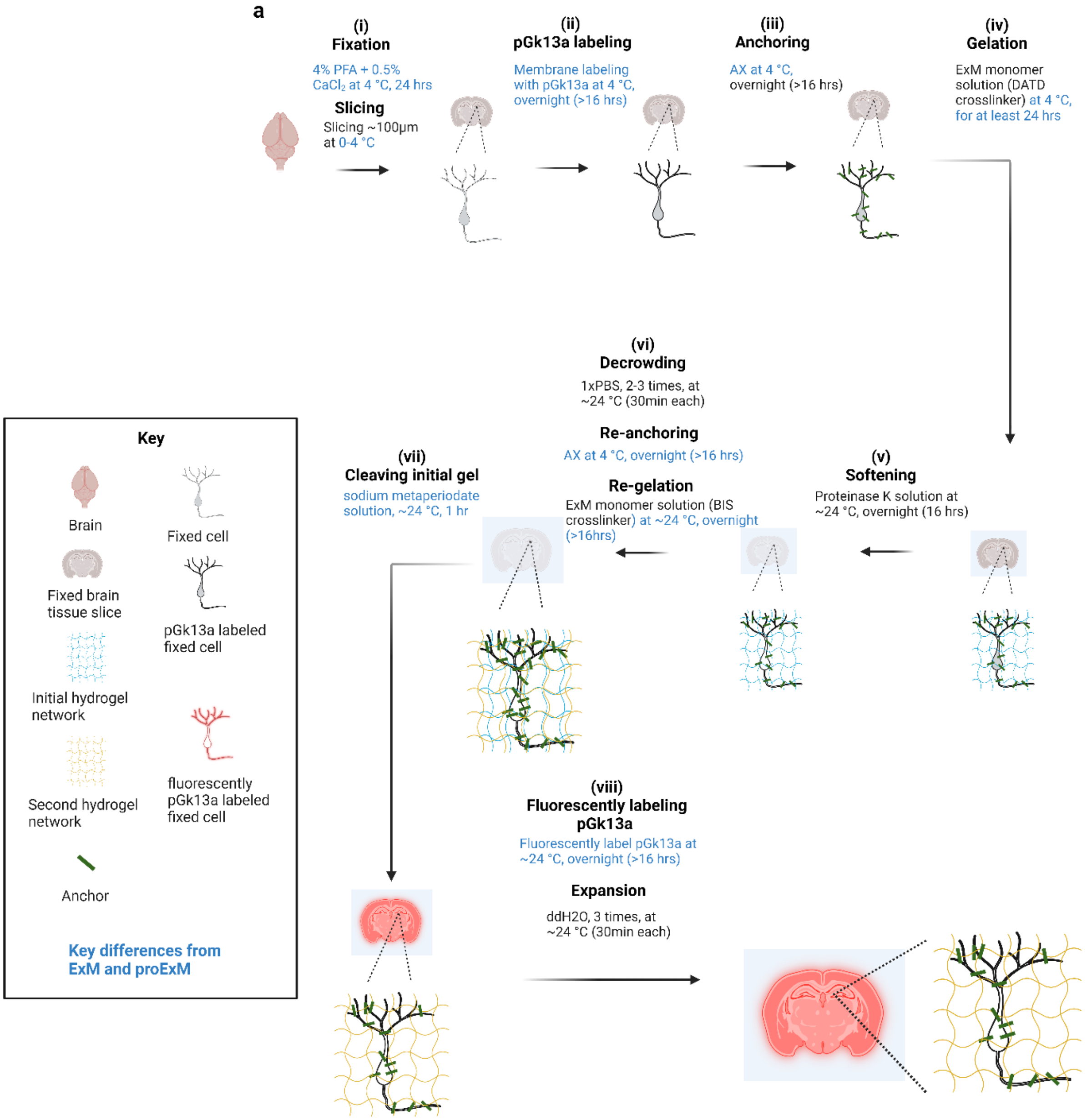

#### Supplementary Figure 21

(**a**) umExM with double gelation workflow schematic. Blue-colored captions highlight the key differences from ExM[3] and proExM[4], whereas black captions highlight steps similar to those in ExM and proExM. (**a.i**) A specimen is chemically fixed with 4% paraformaldehyde (PFA) + 0.5% calcium chloride ($CaCl_2$) at 4 ºC for 24 hours. The brain is sliced on a vibratome at 100 µm thickness at 4 ºC. (**a.ii**) The specimen is treated with pGk13a at 4 ºC overnight (unless otherwise noted, overnight means >16 hours throughout the figure). (**a.iii**) The specimen is treated with acrylic acid N-hydroxysuccinimide ester (AX) at 4 ºC overnight. (**a.iv**) The specimen is embedded in an expandable hydrogel (made with cleavable crosslinker DATD) at 4 ºC for at least 24 hours. (**a.v**) The specimen is mechanically softened with enzymatic cleavage of proteins (i.e., specific cleavage with proteinase-k) at room temperature (~24 ºC), overnight. (**a.vi**) The specimen-embedded hydrogel is treated with PBS to partially expand it. Next, the sample is treated with AX as we did in (**a.iii**). Subsequently, the sample is gelled again but with a monomer solution that contains the non-cleavable crosslinker N,N -Methylenebis(acrylamide) (BIS) at room temperature (~24 ºC), overnight. (**a.vii**) Then the specimen-embedded hydrogel is incubated in a gel cleaving solution (containing sodium metaperiodate) at room temperature (~24 ºC) for 1 hour. This step cleaves the initial gel that was formed in (**a.iv**). (**a.viii**) Finally, the pGk13a, that is anchored to the gel matrix, is fluorescently labeled via click-chemistry (i.e., DBCO-fluorophore) at room temperature, overnight. The specimen-embedded hydrogel is expanded with water at room temperature for 1.5 hours (exchanging water every 30 minutes).

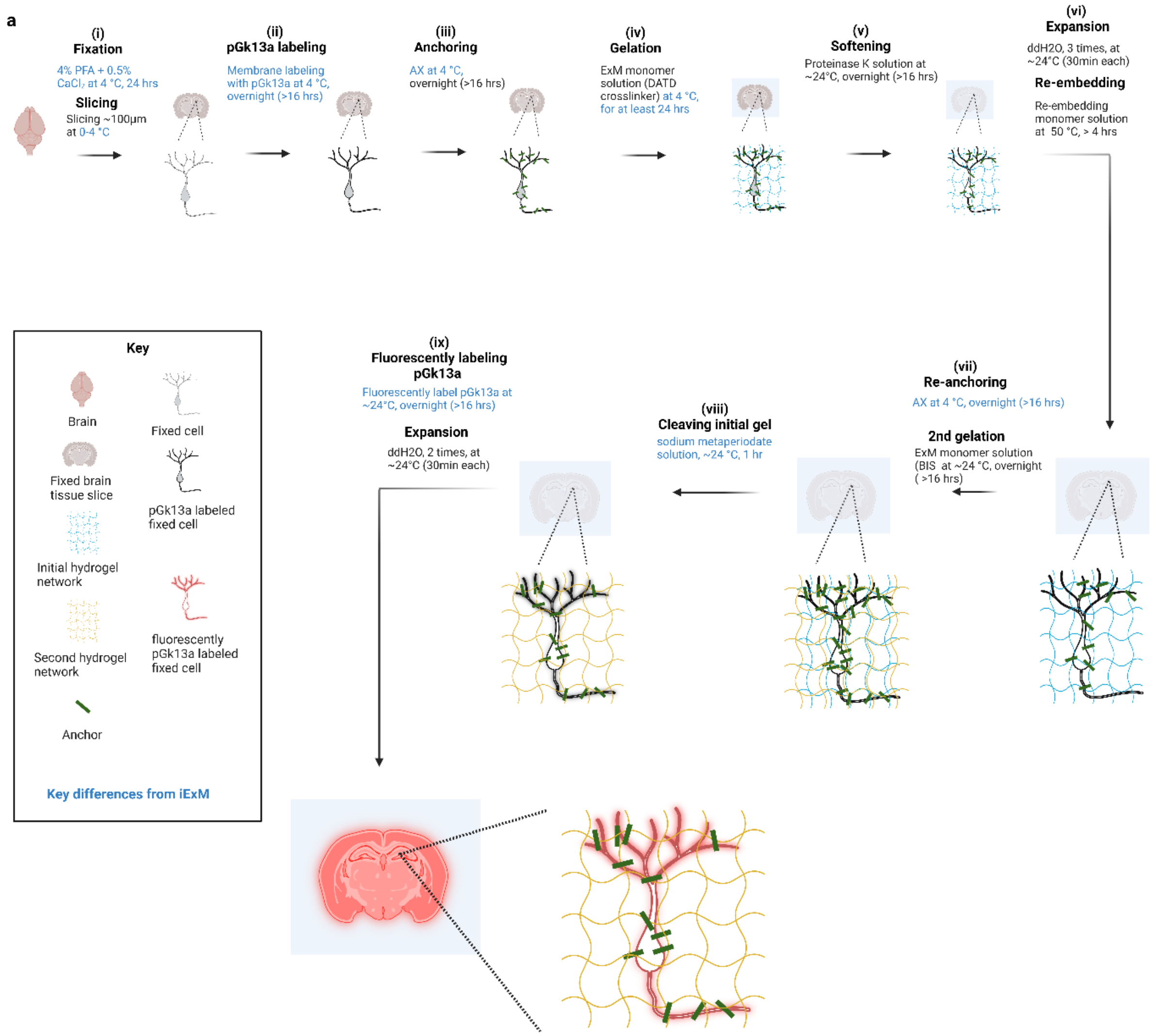

#### Supplementary Figure 22

(**a**) Iterative form of umExM workflow schematic. Blue-colored captions highlight the key differences from iExM[17], whereas black captions highlight steps similar to those in iExM. (**a.i**) A specimen is chemically fixed with 4% paraformaldehyde (PFA) + 0.5% calcium chloride ($CaCl_2$) at 4 ºC for 24 hours. The brain is sliced on a vibratome at 100 µm thickness at 4 ºC. (**a.ii**) The specimen is treated with pGk13a at 4 ºC overnight (unless otherwise noted, overnight means >16 hours throughout the figure). (**a.iii**) The specimen is treated with acrylic acid N-hydroxysuccinimide ester (AX) at 4 ºC overnight. (**a.iv**) The specimen is embedded in an expandable hydrogel (made with cleavable crosslinker DATD[15]) at 4 ºC for at least 24 hours. (**a.v**) The specimen-embedded hydrogel is mechanically softened with enzymatic cleavage of proteins (i.e., specific cleavage with proteinase-k) at room temperature (~24 ºC), overnight. (**a.vi**) The specimen-embedded hydrogel is expanded with water at room temperature for 1.5 hours (exchanging water every 30 minutes). Then, the specimen-embedded hydrogel is re-embedded into a non-expandable hydrogel at 50 ºC, for >4 hours. (**a.vii**) Next, the sample is treated with AX as we did in (**a.iii**). Subsequently, the sample is gelled again but with a monomer solution that contains the non-cleavable crosslinker (made with BIS) at room temperature (~24 ºC), overnight. (**a.viii**) Then the specimen-embedded hydrogel is incubated in a gel cleaving solution (contains sodium metaperiodate) at room temperature (~24 ºC) for 1 hour. This step cleaves the initial gel and re-embedding gel that was formed in (**a.iv**) and (**a.vi**). (**a.ix**) Finally, the pGk13a, that is anchored to the gel matrix, is fluorescently labeled via click-chemistry (i.e., DBCO-fluorophore) at room temperature, overnight. The specimen-embedded hydrogel is expanded with water at room temperature for 1.5 hours (exchanging water every 30 minutes).

### Supplementary Tables

#### Supplementary Table 1

The comparison table excludes TREx[18], LExM[19], TRITON ExM[20], uniExM[21], and sphingolipid ExM[22] as they didn't show membrane or lipid labeling in tissue.

"N/A" denotes not available
* Measured with 100% laser and 100ms exposure time and 60x, 1.27NA water immersion lens

| | ***Protocol*** | **umExM +mExM** | **Magnify** | **pan-ExM-t** | **clickExM** |
|---|---|---|---|---|---|
| **Protocol** | *membrane (lipid) labeling applied to mouse brain tissue?* | Yes | Yes | Yes | Yes |
| **Protocols for mouse brain tissue** | *whether it can be used with conventional fixatives* | Yes | Yes | Yes | No, requires acute brain slice |
| | *which probe?* | pGk13a | DiD | PacSch | alkyne-choline |
| | *cost of the probe? (as of Oct. 2023)* | $150/1mg | $5/1mg | $791/1mg | $15/1mg |
| | *Can clearly visualize plasma membrane?* | Yes | No | No | No |
| | *What other membrane it can visualize?* | -Mitochondria membrane<br>-Nuclear membrane<br>-Ciliary membrane<br>-Extracellular vesicle membrane<br>-Myelin sheaths | -Mitochondria membrane<br>-Bloodvessel membrane<br>-Myelin sheaths | -ER membrane<br>-Mitochondria membrane<br>-Nuclear membrane<br>-Myelin sheaths | unclear |
| | *Claimed ultrastructure is preserved?* | Yes | No | Yes | No |
| | *How they validate ultrastructure preservation?* | Quantitatively comparing the diameter of axon and cilia to previously reported values from EM and STED | N/A | Quantitatively comparing extracellular space content to previously reported values from STED | N/A |
| | *Thickest tissue applied?* | 100µm | 80µm | 100µm | 150µm |
| | *Which brain region?* | Cortex<br>Hippocampus<br>3rd ventricle<br>Choroid plexus<br>Corpus Callosum | unclear | Cortex<br>Hippocampus | Cortex |
| | *signal-to-background of the membrane signal** | ~80 | didn't measured | didn't measured | didn't measured |

| | | | | | |
|---|---|---|---|---|---|
| | *continuity analysis performed?* | Yes | No | No | No |
| | *Post-expansion staining of proteins?* | Yes | Yes | Yes | No |
| | *Post-expansion staining of RNAs?* | Yes | Yes | No | No |
| | *How many antibodies are demonstrated to be used at once?* | 1 | 2 | 2 | 0 |
| | *How many steps involved to reach the highest claimed expansion* | -1 membrane labeling step<br>-2 anchoring steps<br>-3 gelation steps<br>-1 softening step<br>-1 cleaving step<br>-2 expansion step | -1 membrane labeling step<br>-1 anchoring step<br>-1 gelation step<br>-1 softening step<br>-1 expansion step | -1 membrane labeling step<br>-1 anchoring step<br>-3 gelation steps<br>-1 softening step<br>-1 cleaving step<br>-2 expansion steps | -1 membrane labeling step<br>-1 anchoring step<br>-1 gelation step<br>-1 softening step<br>-1 expansion step |
| **Resolution** | *Claimed highest expansion factor* | 12 | 11 | 24 | 4.5 |
| | *Claimed the highest resolution with conventional microscopy* | 35 | didn't measure | didn't measure | didn't measure |
| | *How do they measure the resolution?* | Fourier-ring correlation | N/A | N/A | N/A |
| **Utility** | Segmentation of the cell body? | Yes | No | No | No |
| | Segmentation of the dendrite? | Yes | No | No | No |
| | Segmentation of the axon? | Yes, down to roughly 200nm diameter axons | No | No | No |
| | Tracing of the axon? | Yes | No | No | No |

#### Supplementary Table 2

##### pGk5b stock solution (prepared at RT and immediately stored at -20 ⁰C):

| Reagent | Amount | Final concentration |
|---|---|---|
| pGk5b | 1mg | 10mg/1ml |
| Anhydrous DMSO (Thermo Fisher, cat. no. D12345) | 50µl | |
| Water (Thermo Fisher, cat. no. 10977015) | 50µl | |
| **Total** | **100µl** | |

##### pGk5b membrane labeling stock solution (prepared fresh and used immediately at 4 ⁰C)

| Reagent | Stock concentration | Amount (ml) |
|---|---|---|
| pGk5b stock solution (see above) | 10mg/1ml | 0.01 |
| PBS (Corning, cat. no. 21031CM)* | | 0.99 |
| **Total** | | **1** |

* Chilled on ice before use

##### AcX stock solution (prepared fresh and used immediately at 4 ⁰C*)

| Reagent | Stock concentration | Amount (ml) |
|---|---|---|
| Acryloyl-X SE (Thermo Fisher, cat. no. A20770) | 10mg/ml in DMSO** | 0.01 |
| PBS (Corning, cat. no. 21031CM) | | 0.99 |
| **Total** | | **1** |

* Aliquot 20ul into a PCR tube, and store at -20 ⁰C in a sealed container (e.g., 50mL tube) with drying agents (e.g., Drierite)
** Anhydrous DMSO (Thermo Fisher, cat. no. D12345)

##### Monomer solution aka StockX (9.4ml, aliquoted to 10 tubes of 940µl and stored at -20 ⁰C):

| Reagent | Stock concentration* | Amount (ml) |
|---|---|---|
| Sodium Acrylate (Sigma, cat. no. 408220) | 38 | 2.25 |
| Acrylamide (Sigma, cat. no. A8887) | 50 | 0.5 |
| N,N′-Methylenebisacrylamide (Sigma, cat. no. M7279) | 2 | 0.75 |
| Sodium chloride (Thermo Fisher, cat. no. BP358-212) | 29.2 | 4 |
| 10xPBS (Thermo Fisher, cat. no. 70011044) | 10x | 1 |
| Water (Thermo Fisher, cat. no. 10977015) | | 0.9 |
| **Total** | | **9.4** |

*All concentrations are in g/100 ml except 10xPBS. All stock solutions are formulated in water (Thermo Fisher, cat. no. 10977015).

##### Gelling solution (1ml, prepared at 4 ⁰C, gelled at 37 ⁰C):

| Reagent | Stock concentration* | Amount (µl) |
|---|---|---|
| Monomer Solution (see above) | 1x | 940 |
| 4-hydroxy-TEMPO (Sigma, cat. no. 176141) | 0.5 | 20 |

| | | |
|---|---|---|
| TEMED (Sigma, cat. no. T7024) | 10 | 20 |
| APS (Thermo Fisher, cat. no. 17874) | 10 | 20 |
| **Total** | | **1000** |

*All concentrations are in g/100 ml except Monomer Solution. All stock solutions are formulated in water (Thermo Fisher, cat. no. 10977015).

To make the gelling solution, add 20µl of 4-hydroxy-TEMPO solution (0.005g/ml in water) and 20µl of TEMED solution (0.1g/ml in water) to 940µl of monomer solution, vortex for 2-3 seconds, add 20µl of APS solution (0.1g/ml in water), vortex for 2-3 seconds.

**Digestion buffer* (100ml, prepared and stored at RT, applied at 37 $^{o}$C):**

| **Reagent** | **Stock concentration** | **Amount** |
|---|---|---|
| Tris pH 8.0 (Thermo Fisher, cat. no. AM9856) | 1M | 5ml |
| EDTA (Thermo Fisher, cat. no. 15575020) | 0.5M | 0.2ml |
| Triton X-100 (Sigma, cat. no. X100) | 10% | 5ml |
| NaCl (Sigma, cat. no. S5886) | >99% solid | 5.85g |
| Water (Thermo Fisher, cat. no. 10977015) | | 84ml |
| **Total** | | **100ml** |

To formulate the Digestion solution, dilute Proteinase-K (NEB, cat. no. P8107S) at 1:100 dilution in Digestion buffer. All stock solutions are formulated in water (Thermo Fisher, cat. no. 10977015).

**Fixation Reversal buffer (10ml, prepared at RT and used immediately):**

| **Reagent** | **Stock concentration*** | **Amount** |
|---|---|---|
| PEG20000 (Sigma, cat. no. 95172-250G-F) | 5% | 1ml |
| DTT (Thermo Fisher, cat. no. R0862) | >97% solid | 154.3mg |
| SDS (Thermo Fisher, cat. no. AM9820) | 20% | 2ml |
| Tris pH8 (Thermo Fisher, cat. no. AM9856) | 1M | 1ml |
| Water (Thermo Fisher, cat. no. 10977015) | | 5.9ml |
| **Total** | | **10ml** |

*****All stock solutions are formulated in water (Thermo Fisher, cat. no. 10977015).

**List of antibodies used for mExM:**

| **Antigen** | **Species** | **Company** | **Catalog no** | **Note** |
|---|---|---|---|---|
| **Tom20** | Rabbit | CST* | 42406S | Identified as Vendor1 in Supp. Fig. 5d, mitochondria bar |
| **Tom20** | Mouse | SCBT** | sc-17764 | Identified as Vendor2 in Supp. Fig. 5d, mitochondria bar |
| **Nup98** | Rabbit | CST* | 2597S | |
| **MBP** | Rabbit | CST* | 78896S | Identified as Vendor1 in Supp. Fig. 5d, myelin bar |
| **MBP** | Rabbit | Abcam | ab40390 | Identified as Vendor2 in Supp. Fig. 5d, myelin bar |
| **MBP** | Chicken | AVES | AB_2313550 | Identified as Vendor3 in Supp. Fig. 5d, myelin bar |

* Cell Signaling Technology
**Santa Cruz Biotechnology

#### Supplementary Table 3

##### Fixative solution (prepared fresh and used immediately):

| Reagent | Stock concentration | Amount |
|---|---|---|
| PFA (Electron Microscopy Sciences, cat. no. 15710) | 16% | 10ml |
| $CaCl_2$ (Millipore Sigma, cat. no. C4901) | ≥97% solid | 0.2g |
| Sodium cacodylate buffer, pH 7.4 (Electron Microscopy Sciences, cat. no. 11653)* | 200mM | 30ml |
| **Total**** | | 40ml |

*Chilled on ice before use
**Kept on ice during perfusion

##### pGk13a stock solution (prepared at RT and immediately stored at -20 $^0$C):

| Reagent | Amount | Final concentration |
|---|---|---|
| pGk13a | 1mg | 11.1mg/1ml |
| Anhydrous DMSO (Thermo Fisher, cat. no. D12345) | 45µl | |
| Water (Thermo Fisher, cat. no. 10977015) | 45µl | |
| **Total** | **90µl** | |

##### pGk13a membrane labeling stock solution (prepared fresh and used immediately at 4 $^0$C)

| Reagent | Stock concentration | Amount (ml) |
|---|---|---|
| pGk13a stock solution (see above) | 11.1mg/ml | 0.015 |
| PBS (Corning, cat. no. 21031CM)* | 1x | 0.985 |
| **Total** | | **1** |

* Chilled on ice before use

##### AX buffer solution (prepared fresh and stored at 4 $^0$C)

| Reagent | Amount | Final concentration |
|---|---|---|
| MES (Sigma, cat. no. M3058) | 0.434mg | 0.4344g/20ml |
| 5M Sodium chloride (Thermo Fisher, cat. no. BP358-212) | 0.6ml | |
| 1M HCl | 1ml | |
| Water (Thermo Fisher, cat. no. 10977015)* | 18.4ml | |
| **Total** | 20ml | |

* Chilled on ice before use

##### AX stock solution (prepared fresh and used immediately at 4 $^0$C, and stored at -20 $^0$C*)

| Reagent | Stock concentration | Amount (ml) |
|---|---|---|
| Acrylic acid N-hydroxysuccinimide ester (Thermo Fisher, cat. no. AC400300010) | 10mg/ml in DMSO** | 0.006 |

| | | |
|---|---|---|
| AX buffer solution (see above) | | 0.994 |
| **Total** | | **1** |

* Aliquot 20ul into a PCR tube, and store at -20 ºC in a sealed container (e.g., 50mL tube) with drying agents (e.g., Drierite)
** Anhydrous DMSO (Thermo Fisher, cat. no. D12345)

**umExM monomer solution (9.4ml, aliquoted to 10 tubes of 940µl and stored at -20 ºC):**

| Reagent | Stock concentration* | Amount (ml) |
|---|---|---|
| Sodium Acrylate (Sigma, cat. no. 408220) | 38 | 2.25 |
| Acrylamide (Thermo Fisher, cat. no. 15512023) | 50 | 0.5 |
| N,N'-Diallyl-L-tartardiamide (Alfa Aesar, cat. no. A12195-30) | 9 | 0.75 |
| 5M Sodium chloride (Thermo Fisher, cat. no. BP358-212) | 29.2 | 4 |
| 10xPBS (Thermo Fisher, cat. no. 70011044) | 10x | 1 |
| Water (Thermo Fisher, cat. no. 10977015) | | 0.9 |
| **Total** | | **9.4** |

*All concentrations are in g/100 ml except 10xPBS.

**umExM gelling solution (1ml, prepared and gelled at 4 ºC):**

| Reagent | Stock concentration* | Amount (µl) |
|---|---|---|
| umExM monomer Solution (see above) | 1x | 940 |
| 4-hydroxy-TEMPO (Sigma, cat. no. 176141) | 0.5 | 20 |
| TEMED (Sigma, cat. no. T7024) | 10 | 20 |
| APS (Thermo Fisher, cat. no. 17874) | 10 | 20 |
| HCl | 1M | 14 |
| **Total** | | **1014**** |

*All concentrations are in g/100 ml except umExM Monomer Solution. All stock solutions are formulated in water (Thermo Fisher, cat. no. 10977015).
**To make umExM gelling solution, add 20µl of 4-hydroxy-TEMPO solution (0.005g/ml in water) and 20µl of TEMED solution (0.1g/ml in water) to 940µl of umExM monomer Solution, vortex for 2-3 seconds, add 20µl of APS solution (0.1g/ml in water), vortex for 2-3 seconds, and add 4µl of 1M HCl, and vortex for 2-3 second.

**umExM Digestion buffer* (100ml, prepared and applied at RT, and stored at 4 ºC):**

| Reagent | Stock concentration | Amount |
|---|---|---|
| Tris pH 8.0 (Thermo Fisher, cat. no. AM9856) | 1M | 5ml |
| EDTA (Thermo Fisher, cat. no. 15575020) | 0.5M | 0.2ml |
| Saponin (Sigma, cat. no. 84510) | 10% | 5ml |
| 5M Sodium chloride (Thermo Fisher, cat. no. BP358-212) | 5M | |
| Water (Thermo Fisher, cat. no. 10977015) | | 84ml |
| **Total** | | **100ml** |

*To formulate the Digestion solution, dilute Proteinase-K (NEB, cat. no. P8107S) at 1:100 dilution in Digestion buffer. All stock solutions are formulated in water (Thermo Fisher, cat. no. 10977015).

**Trypsin+Lys-C softening solution (2ml, prepared fresh and applied at RT):**

| Reagent | Amount | Final concentration |
|---|---|---|
| Trypsin/Lys-C (Thermo Fisher, cat. no. A40007) | 20µg | 20µg/2ml |
| TRIS (1M), pH 8.0 (Thermo Fisher, cat. no. AM9855G) | 0.1ml | |
| PBS (Corning, cat. no. 21031CM) | 1.9ml | |

| | |
|---|---|
| **Total** | **2ml** |

**Second monomer solution (34.5ml, prepared fresh):**

| **Reagent** | **Stock concentration*** | **Amount (ml)** |
|---|---|---|
| Acrylamide (Thermo Fisher, cat. no. 15512023) | 50 | 10 |
| N,N'-Diallyl-L-tartardiamide (Alfa Aesar, cat. no. A12195-30) | 5 | 5 |
| Water (Thermo Fisher, cat. no. 10977015) | | 0.9 |
| **Total** | | **34.5** |

*All concentrations are in g/100 ml.

**Second gelling solution (50ml, prepared fresh):**

| **Reagent** | **Stock concentration*** | **Amount (ml)** |
|---|---|---|
| Second monomer solution | 1x | 34.5 |
| N,N'-Diallyl-L-tartardiamide (Alfa Aesar, cat. no. A12195-30) | 5 | 5 |
| TEMED (Sigma, cat. no. T7024) | 10 | 0.25 |
| APS (Thermo Fisher, cat. no. 17874) | 10 | 0.25 |
| **Total** | | **50** |

*All concentrations are in g/100 ml except cleavable second monomer solution.

To make the second gelling solution, 250μl of TEMED solution (0.1g/ml in water) to 34.5mL of second monomer solution, vortex for ~10 seconds, add 250μl of APS solution (0.1g/ml in water), vortex for ~10 seconds.

**Third monomer solution (9.4ml, aliquoted to 10 tubes of 940μl and stored at -20 $^0$C):**

| **Reagent** | **Stock concentration*** | **Amount (ml)** |
|---|---|---|
| Sodium Acrylate (Sigma, cat. no. 408220) | 38 | 2.25 |
| Acrylamide (Sigma, cat. no. A8887) | 50 | 0.5 |
| N,N′-Methylenebisacrylamide (Sigma, cat. no. M7279) | 2 | 1 |
| Sodium chloride (Thermo Fisher, cat. no. BP358-212) | 29.2 | 4 |
| 10xPBS (Thermo Fisher, cat. no. 70011044) | 10x | 1 |
| Water (Thermo Fisher, cat. no. 10977015) | | 0.65 |
| **Total** | | **9.4** |

*All concentrations are in g/100 ml except 10xPBS.

**Third gelling solution (prepared in fresh and applied at RT):**

| **Reagent** | **Stock concentration*** | **Amount (μl)** |
|---|---|---|
| umExM monomer Solution (see above) | 1x | 940 |
| 4-hydroxy-TEMPO (Sigma, cat. no. 176141) | 0.5 | 20 |
| TEMED (Sigma, cat. no. T7024) | 10 | 20 |
| APS (Thermo Fisher, cat. no. 17874) | 10 | 20 |
| **Total** | | **1000**** |

*All concentrations are in g/100 ml.
** To make the third gelling solution, add 20μl of 4-hydroxy-TEMPO solution (0.005g/ml in water) and 20μl of TEMED solution (0.1g/ml in water) to 940μl of monomer solution, vortex for 2-3 seconds, add 20μl of APS solution (0.1g/ml in water), vortex for 2-3 seconds.

**List of antibodies used for umExM:**

| Antigen | Species | Company | Catalog no |
|---|---|---|---|
| **SV2A** | Rabbit | Abcam | 50-194-3924 |
| **PSD95** | Rabbit | Thermo Fisher | MA1-046 |

### Supplementary Methods

#### Electron microscope imaging, visualization, and analysis

To validate pGk5b labeling by electron microscopy (EM), tissue slices of 100 μm thickness were treated with lipid labeling solution as we used for mExM, except we used azide instead of biotin as the linkable group of the lipid stain (pGk5a for short): we first incubated the tissue in 1ml of pGk5 (0.1μg/ml in ice-cold PBS) at 4 ºC overnight (>16hrs) to let the labels diffuse and intercalate thoroughly throughout. Subsequently, the sample was washed 2x using PBS at 4 ºC for 1 hour each to remove any excess lipid label. Then the sample was placed in 2% PFA, 2% glutaraldehyde in PBS at 4 ºC for 6 hours for post-fixation of the sample for EM staining. This fix also served for further EM processing, to preserve the state of ultrastructure[23,24]. These steps were performed at 4 ºC to promote the stability of the lipids and the lipid label in the sample. The sample was moved to 1.8nm undecagold-DBCO conjugate solution (2.5mg/1mL Nanopartz, part no. CK11) at 4 ºC for 12 hours. The azide-DBCO chemistry served to link the lipid label with a gold nanoparticle. Thereafter the sample was washed 3x with 0.15M sodium cacodylate buffer at room temperature for 30 minutes each to remove unbound nanoparticles. The samples were sent to the Harvard Medical School Electron Microscopy Core to be stained, then embedded and sliced using a standard EM preparation protocol[23]. In summary, the tissue was stained with 1% uranyl acetate (UA) for 1 hour at room temperature, embedded in resin, and sliced in ultrathin sections (40nm thickness). As discussed in **Supp. Fig. 1**, we decided to use a common UA staining protocol (1% UA for 1 hour at RT) to enhance the pGk5a signals on top of signals from gold nanoparticles as UA can react to amino groups of pGk5a. As for control experiments, the protocol was adjusted by replacing pGk5a staining with common osmium staining (1% $OsO_4$ for

1 hour at RT). Samples were imaged on a JEOL 1200EX transmission electron microscope using 80keV transmitted voltage. The images were captured with an AMT 2k CCD camera. Acquired images were processed with a Gaussian filter (Radius = 1) and Enhance contrast (Saturated pixels = 10.5%) function in ImageJ (version 1.53q). pGk5a treated sample without $OsO_4$ clearly showed membranes (e.g., mitochondrial membrane and vesicle membranes; **Supp. Fig. 1a**) similar to the control experiment (i.e., that is, only with $OsO_4$; **Supp. Fig. 1b**) but with slightly lower contrast.

**Cell preparation**

We first inserted a 13-mm-diameter coverslip (Thermo Fisher, catalog no. 174950) into one well of a 24-glass well plate (Cellvis 24 WELL GLASS BTTM PLATE 20/CS, catalog no. NC0397150). Then, either HEK293 or HeLa or U2OS cells were plated in the well (~40k cells/ml in cell culture medium (described in next paragraph) per well.) The plate was then moved to a humidified cell culture incubator (set at 37°C, 20% oxygen, and 5% CO2) for at least 6 hours for cells to adhere. The cells were fixed with 4% paraformaldehyde (PFA) and 0.1% glutaraldehyde in Dulbecco's 1x phosphate buffered saline (PBS) at room temperature (RT) for 15 minutes. Fixed cells were washed 4 times with PBS for 10 minutes each at 4 $^{o}$C, and kept in PBS at 4 $^{o}$C.

For HEK293 cell culture medium, we used Dulbecco’s modified Eagle’s medium (DMEM, Corning, catalog no. 10013CV) supplemented with 10% heat-inactivated fetal bovine serum (FBS, Thermo Fisher, catalog no. A3840001), 2 mM GlutaMax (Thermo Fisher Scientific, cat. No. 3505006), and 1% penicillin-streptomycin (Thermo Fisher, catalog no. 15140122). For HeLa cell culture medium, we used DMEM supplemented with 10% heat-inactivated fetal bovine serum (FBS) and 1% penicillin-streptomycin (Thermo Fisher, catalog no. 15140122). For U2OS cell culture medium, we used Dulbecco's modified Eagle's medium (DMEM, Corning, catalog no. 10013CV) supplemented with 10% heat-inactivated fetal bovine serum (FBS, Thermo Fisher, catalog no. A3840001), 1% penicillin-streptomycin (Thermo Fisher, catalog no. 15140122) and 1% sodium pyruvate (Thermo Fisher, catalog no. 11360070).

#### Transduction of cells via BacMam virus

The adherent cells were prepared as described in the Cell Preparation section. The cells were transduced by directly adding 12µl of BacMam reagent (either CellLight™ Mitochondria-GFP, catalog no. C10508 or CellLight™ ER-GFP, catalog no. C10590) to the cell medium. The cells were then placed in the culture incubator overnight (>16hrs). The cells were then fixed and washed as described in **the Cell Preparation section** in **Supplementary Methods**.

#### Brain tissue preparation for mExM

All procedures involving animals were in accordance with the US National Institutes of Health Guide for the Care and Use of Laboratory Animals and approved by the Massachusetts Institute of Technology Committee on Animal Care. Wild type (both male and female, C57BL/6 or Thy1-YFP, 6-8 weeks old, from either Taconic or JAX) mice were first terminally anesthetized with isoflurane. Then, ice-cold PBS was transcardially perfused until the blood cleared (approximately 25ml). For all mExM experiments, the mice were then transcardially perfused with 4% paraformaldehyde (PFA) and 0.1% glutaraldehyde in ice-cold PBS. The fixative was kept on ice during perfusion. After the perfusion step, brains were dissected out, stored in fixative at 4 $^{o}$C for 12 hours for further fixation, and sliced on a vibratome (Leica VT1000S) at 100 μm thickness. For the slicing, the tray was filled with ice-cold PBS, and the tray was surrounded by ice. The slices were kept in PBS at 4 ºC overnight for washing and storing.

**mExM for cells**

1. The fixed cells (as described in the **Cell Preparation** section in **Supplementary Methods**) were incubated in the pGk5b solution (**Supp. Table 2,** "pGk5b membrane labeling stock solution") at 4 ºC overnight.
2. The cells were then incubated in the AcX solution (**Supp. Table 2,** "AcX stock solution") overnight at 4 ºC. Then we washed with ice-cold PBS 2 times, 30min each at 4 ºC.
3. The cells were then incubated in the gelling solution (**Supp. Table 2, "**Gelling solution") 30min at 4 ºC for pre-gelation. During this step, the gelation chamber was constructed as described previously[3]. In summary, we placed two spacers (VWR, catalog no. 48368-085) on a microscope slide (VWR, catalog no. 48300-026). The spacers were separated from each other enough so that an adherent cell-containing cover glass could be placed in between them. The adherent cell-containing cover glass was then placed between the spacers on the slide. We then placed the lid (VWR, catalog no. 87001-918) on top of the spacers, covering the cell-containing cover glass. We then fully filled the empty space between the cells and spacers with the gelling solution. Next, the chamber was transferred to a 37 ºC incubator to initiate free-radical polymerization. After 2 hours, the gelation chamber containing cells was taken out
4. The gel was trimmed with a razor blade (VWR, cat. no. 55411-050) and transferred from the chamber to a 6-well plate (Thermo Fisher, catalog no. 140675) that contained proteinase K digestion buffer (**Supp. Table 2, "**Digestion buffer") in the well (3mL of

digestion buffer per well). The gel was then digested at 37C on a shaker overnight (> 16 hours). After digestion, the gel was washed 4 times in PBS at RT, 30 minutes each.

5. The digested gels were labeled with 0.3mg/ml of streptavidin labeled with Atto 565 (Atto 565-Streptavidin; Sigma Aldrich, catalog no. 56304-1MG-F) buffered in PBS overnight at RT, and then washed 4 times in PBS at room temperature (RT), 30 minutes each.
6. The gels were placed 4 times in excess water at RT for expansion, 30 minutes each.

**mExM for brain tissue slices**

1. The fixed tissue slices (as described in the **Brain tissue preparation for mExM** section in **Supplementary Methods**) were incubated in a lipid labeling solution (**Supp. Table 2,** “Lipid labeling stock solution”) at 4 $^{o}$C overnight (>16 hours) to let the labels diffuse and intercalate thoroughly throughout the tissue slices.
2. The tissue slices were then incubated in an AcX stock solution (**Supp. Table 2,** “AcX stock solution”) overnight (>16 hours) at 4 $^{o}$C. The tissue was then washed 2 times in PBS at 4 $^{o}$C, 1 hour each.
3. The tissue slices were then incubated in gelling solution (**Supp. Table 2, “**Gelling solution”) 30min at 4 $^{o}$C for pre-gelation incubation. During this step, the gelation chamber was constructed as previously described[3]. In summary, we placed two spacers (VWR, catalog no. 48368-085) on a microscope slide (VWR, catalog no. 48300-026). The two spacers were separated from each other enough so that the brain tissue slice could be placed in between them. The brain tissue slice was placed between the spacers. We then placed the lid (VWR, catalog no. 87001-918) on top of the spacers as well as the brain tissue slice. We then fully filled the empty space between the brain tissue slice and

spacers with the gelling solution. The chamber was transferred to a 37 ºC incubator to initiate free-radical polymerization. After 2 hours, the gelation chamber containing the tissue was taken out.

4. The gel was trimmed with a razor blade (VWR, cat. no. 55411-050) and transferred from the chamber to a 6-well plate (Thermo Fisher, catalog no. 140675) that contained proteinase K digestion buffer (**Supp. Table 2**, “Digestion buffer”) in the well (4mL of digestion buffer per well). The gel was then digested at 37C on a shaker overnight (>16 hours). After digestion, the gel was washed 4 times in PBS at RT, 30 minutes each.
5. The digested gels were labeled with 0.3mg/ml of streptavidin labeled with Atto 565 (Atto 565-Streptavidin; Sigma Aldrich, catalog no. 56304-1MG-F) buffered in PBS overnight at RT, and then washed 4 times in PBS at room temperature (RT), 30 minutes each.
6. The gels were placed 4 times in excess water at RT for expansion, 30 minutes each.

**Immunohistochemistry-compatible mExM**

The aforementioned mExM steps were carried out the same, except for the digestion step (i.e., step4 in the mExM for cells and mExM for brain tissue slices sections). Instead of using the proteinase K digestion buffer, the sample was heated in fixation reversal (FR; Supp Table 1, “Fixation Reversal buffer”) buffer for 30 minutes at 100 ºC and then held for 2 hours at 80 ºC. The FR buffer consisted of 0.5% PEG20000, 100mM DTT, 4% SDS, in 100mM Tris pH8. After this, the FR-digested sample was washed in 1x PBS 4 times at RT for 1 hour before proceeding to the immunohistochemistry steps. The expanded gels were first blocked with MAXblock Blocking Medium (Active Motif, catalog no. 15252) for 4-6 hours at room temperature and incubated in MAXbind Staining Medium (Active Motif, catalog no. 15251) containing primary antibodies at a concentration of 10 μg/ml overnight at 4 ºC. Then, the sample was washed with MAXwash Washing Medium (Active Motif, catalog no. 15254) at RT 4 times, 30 minutes each and subsequently incubated in secondary antibodies buffered in MAXbind Staining Medium at a concentration of 10 μg/ml for 10-12 hours at 4 ºC. Finally, the secondary antibodies were washed, again, with MAXwash Washing Medium at RT 4 times, 30 minutes each time. For primary antibodies, anti-TOM20 (Cell Signaling Technology, catalog no. 42406S, rabbit; Santa Cruz Biotechnology, catalog no. sc-17764, mouse), anti-NUP98 (Cell Signaling Technology, catalog no. 2597S, rabbit), anti-myelin basic protein (MBP; Cell Signaling Technology, catalog no. 78896S, rabbit; Abcam, catalog no. ab40390, rabbit; AVES, catalog no. AB_2313550, chicken), were used. For secondary antibodies, anti-chicken Alexa Fluor Plus 488 (Thermo Fisher, catalog no. A32931), anti-rabbit Alexa Fluor Plus 488 (Thermo Fisher, catalog no. A32731), and anti-mouse Alexa Fluor Plus 647 (Thermo Fisher, catalog no. A32728) were used. After antibody staining, the pGk5b probes that were conjugated to the gel were then labeled with 0.3mg/ml of streptavidin labeled with Atto 565 (Atto 565-Streptavidin; Sigma Aldrich, catalog no. 56304-1MG-F) buffered in PBS overnight at RT, and then washed 4 times in PBS at RT, 30 minutes each. Finally, the gel was placed 4 times in excess water at RT for expansion, 30 minutes each.

**Antibody staining of fluorescent proteins for mExM**

The expanded samples, after either proteinase-k digestion or high-temperature softening, were incubated with MAXblock Blocking Medium (Active Motif, catalog no. 15252) for 4-6 hours at room temperature and incubated in MAXbind Staining Medium (Active Motif, catalog no. 15251) containing fluorophore-conjugated primary antibody against the green fluorescent protein

(GFP) at a concentration of 10 μg/mL overnight (> 16 hours) at 4 ºC. For the primary antibody, we used anti-GFP (Thermo, catalog no. A-21311). Next, the sample was washed with MAXwash Washing Medium (Active Motif, catalog no. 15254) at RT 4 times, 30 minutes each. After antibody staining, the lipid labels that were conjugated to the gel were then labeled with 0.3mg/ml of streptavidin labeled with Atto 565 (Atto 565-Streptavidin; Sigma Aldrich, catalog no. 56304-1MG-F) buffered in PBS overnight at RT, and then washed 4 times in PBS at RT, 30 minutes each. Finally, the gel was placed 4 times in excess water at RT for expansion, 30 minutes each.

**Expansion factor and degree of isotropy analysis**

For umExM, expansion factor and degree of isotropy analysis were carried out with cells. mExM protocol for cells is described **mExM for cells** section in supplementary methods. umExM protocol for cells is similar as mExM for cells, but using the pGk13a membrane labeling stock solution for step 1, AX stock solution for step 2, umExM gelling solution for step 3, umExM Digestion buffer for step 4, and pGk13a was fluorescently labeled with 1ml of Cy3 conjugated DBCO (Cy3 DBCO Click chemistry tools, catalog no. A140-1) buffered in PBS at a concentration of 0.03mg/ml on the shaker (50 rpm) at RT, overnight (>16 hours). The samples were expanded with water.

We evaluated the expansion factor as previously described[25,26]. In particular, we used HEK293 and U2OS cells transfected with BacMam viruses expressing GFP proteins targeted to the matrix of mitochondria. We randomly chose two landmarks in pre-expansion images and found the corresponding landmarks in expanded-cell images, and calculated the ratio.

We evaluated distortion as previously described[25,26]. In summary, we used BacMam virus to express GFP proteins in the matrix of mitochondria in HEK293 and U2OS cells. We imaged the cell with SIM before expansion, and re-imaged the same region after expansion with a confocal microscope. We non-rigidly registered the SIM image and the confocal image, then calculated the root-mean-square (RMS) length measurement error as a function of measurement length for SIM vs. expanded-cell images.

For the iterative form of umExM, expansion factor was measured with slices of fixed mouse brain. We measured the gel size before and after expansion (i.e., after 2nd round of expansion) and divided the measured gel size to obtain expansion factor.

**Colocalization analysis for mExM images**

We performed a colocalization analysis for mExM by adopting recommended colocalization methods for light microscopy studies[27,28]. We first segmented the foreground and background fluorescence of GFP channels of mExM images using the Otsu image processing algorithm[29] as we previously did for segmenting signals[5,30]. We then created a binary signal mask based on the foreground signals and used the signal mask to segment the pGk5b signals. Finally, we evaluated the fraction of expressed GFP and antibody signals that had pGk5b signals by counting the pixels containing pGk5b signals that were above 1x standard deviation below the mean of the pGk5b signal intensity in the image. The analysis was performed with RStudio 2021.09.2+382 with R version 4.1.2.
